# Disease-causing mutations in the G protein β5 β-propeller disrupt its chaperonin-mediated folding trajectory

**DOI:** 10.1101/2025.05.28.656654

**Authors:** Mikaila I. Sass, Deirdre C. Mack, Riley A. Nickles, Caelen A. Jones, Jesus N. Ramirez Torres, Aaron M. Wilson, Rex L. Brunsdale, Cecilia S. Sanders, Samuel L. Cottam, Liam J. Moss, Stefano Maggi, Peter S. Shen, Barry M. Willardson

**Affiliations:** Department of Chemistry and Biochemistry, Brigham Young University, C100 BNSN, Provo, UT 84602, USA; Department of Biochemistry, School of Medicine, University of Utah, 15 N. Medical Drive East, Salt Lake City, UT 84112, USA

**Keywords:** G protein, missense mutations, chaperonin, protein folding, cryo-electron microscopy

## Abstract

The Chaperonin Containing Tailless polypeptide 1 (CCT or TRiC) is an essential cytosolic chaperone that folds multiple protein substrates, including many with β-propeller folds. One β-propeller substrate is the G protein β_5_ subunit (Gβ_5_) of Regulator of G protein Signaling (RGS) complexes that determine the duration of G protein signals in neurons. In recent work, we used cryo-electron microscopy (cryo-EM) to visualize the complete CCT-mediated folding trajectory for Gβ_5_, from an initiating electrostatic interaction of a single β-strand in Gβ_5_ with the CCT5 subunit to a completely folded β-propeller structure. Here, we employed biochemistry and cryo-EM to determine key interactions with CCT that initiate Gβ_5_ folding and how missense mutations in Gβ_5_ that cause severe neurological diseases alter the Gβ_5_ folding trajectory and lead to incompletely folded, trapped intermediates. These findings highlight how CCT recognizes folding substrates, how defects in chaperonin-mediated folding contribute to disease, and how strategies might be designed to stabilize misfolded proteins to restore function.

**Significance:** Electrostatic interactions between the CCT chaperonin and its protein substrates initiate the folding process. Using cryo-EM structure determinations, we found a striking specificity for these interactions that allow CCT-mediated Gβ_5_ folding. Moreover, certain missense mutations in Gβ_5_ lead to misfolding and are associated with neurological disorders. We tracked how these mutations disrupt the normal folding of Gβ_5_ by CCT. Although mutant Gβ_5_ still binds the complex, folding stalls mid-process, leaving the protein trapped in partially folded, non-functional states. These defects arise from disrupted packing of the Gβ_5_ core that interferes with formation of the native structure. Our findings reveal a molecular basis for Gβ_5_ misfolding in disease and suggest pharmacological chaperones that stabilize the folded state might restore proper function.

## Introduction

G protein-coupled receptors (GPCRs) transduce a myriad of extracellular signals across the cell membrane into the cytosol to elicit physiological responses (1–3). These seven transmembrane helical structures bind extracellular ligands, inducing conformational changes that propagate to the inner surface of the plasma membrane where the GPCR interacts with the G protein heterotrimer (composed of Gα, Gβ, and Gγ subunits) to trigger the exchange of guanosine diphosphate (GDP) for guanosine triphosphate (GTP) on the Gα subunit (1, 4, 5). This nucleotide exchange disrupts the interaction of Gα with both the GPCR and the Gβγ dimer, allowing Gα-GTP and Gβγ to regulate downstream effectors (6–9). Gα hydrolysis of GTP to GDP inactivates Gα and allows it to reassociate with Gβγ. However, many Gα isoforms hydrolyze GTP at a rate too slow for rapid physiological responses and require interactions with Regulator of G Signaling (RGS) proteins to accelerate this process (10, 11). Among RGS proteins, members of the R7 family (RGS6,7,9 and 11) associate with Gβ_5_, a unique isoform of Gβ found predominantly in neurons (12), to form a dimer that stabilizes the R7 RGS protein, allowing it to interact with Gα, increase Gα GTPase activity, and thereby set the duration of the G protein signal (13–15).

To perform its function, Gβ_5_ must be folded and assembled into its RGS-Gβ_5_ dimer by molecular chaperones. The primary chaperone for all Gβ subunits is the cytosolic Chaperonin Containing Tailless polypeptide 1 (CCT, also known as the TCP-1 ring complex (TRiC)) and its co-chaperone phosducin-like protein 1 (PhLP1) (16–18). CCT is comprised of two rings with eight paralogous subunits per ring. The rings stack together to create a barrel-like structure with a central chamber where nascent or unfolded proteins bind and are folded (19–21). Each CCT subunit binds and hydrolyzes ATP, which in turn drives a conformation change in CCT that closes the ends of the barrel (22–26). This conformational change traps the client protein to be folded (called the substrate) inside the chamber where it folds. After ATP hydrolysis and phosphate release, the CCT subunits relax back into their open conformation (27), and the folded substrate can dissociate from CCT to interact with its binding partners.

Gβ_5_ and the other four Gβ proteins are part of a structural class of WD40 repeat proteins that adopt a seven-bladed β-propeller fold. The blades are made up of 4-stranded, anti-parallel β-sheets that create a circular structure resembling an airplane propeller (28, 29). To close the β-propeller structure, the last three C-terminal β-strands interact with the first N-terminal β-strand to create the seventh β-sheet (Fig. 1A) (28–30). Many β-propeller proteins rely on CCT to bring these N- and C-terminal strands together (16, 31–36). Using cryo-electron microscopy (cryo-EM), we recently reported the folding trajectory of Gβ_5_ while bound to CCT, demonstrating how CCT orchestrates the folding of a β-propeller protein (37). Remarkably, CCT promotes Gβ_5_ folding through primarily hydrophilic interactions between surface residues of the β-propeller blades with the apical domains of CCT that leave the hydrophobic core of Gβ_5_ free to coalesce into its β-propeller structure. Folding initiates in the middle of the β-propeller through an electrostatic interaction between R269 (Gβ_5_ long splice form amino acid numbering) of blade 4 and E256 of the CCT5 apical domain. Folding then propagates around the β-propeller with blades 3 and 5 making hydrophilic interactions with CCT2 and CCT7, the subunits adjacent to CCT5 in the ring, respectively. The process continues in both directions around the rest of the β-propeller without any additional direct interactions with CCT. Blade 1 is the last to fold and its interaction with blade 2 closes the β-propeller structure (Fig. 1B).

**Figure 1.**
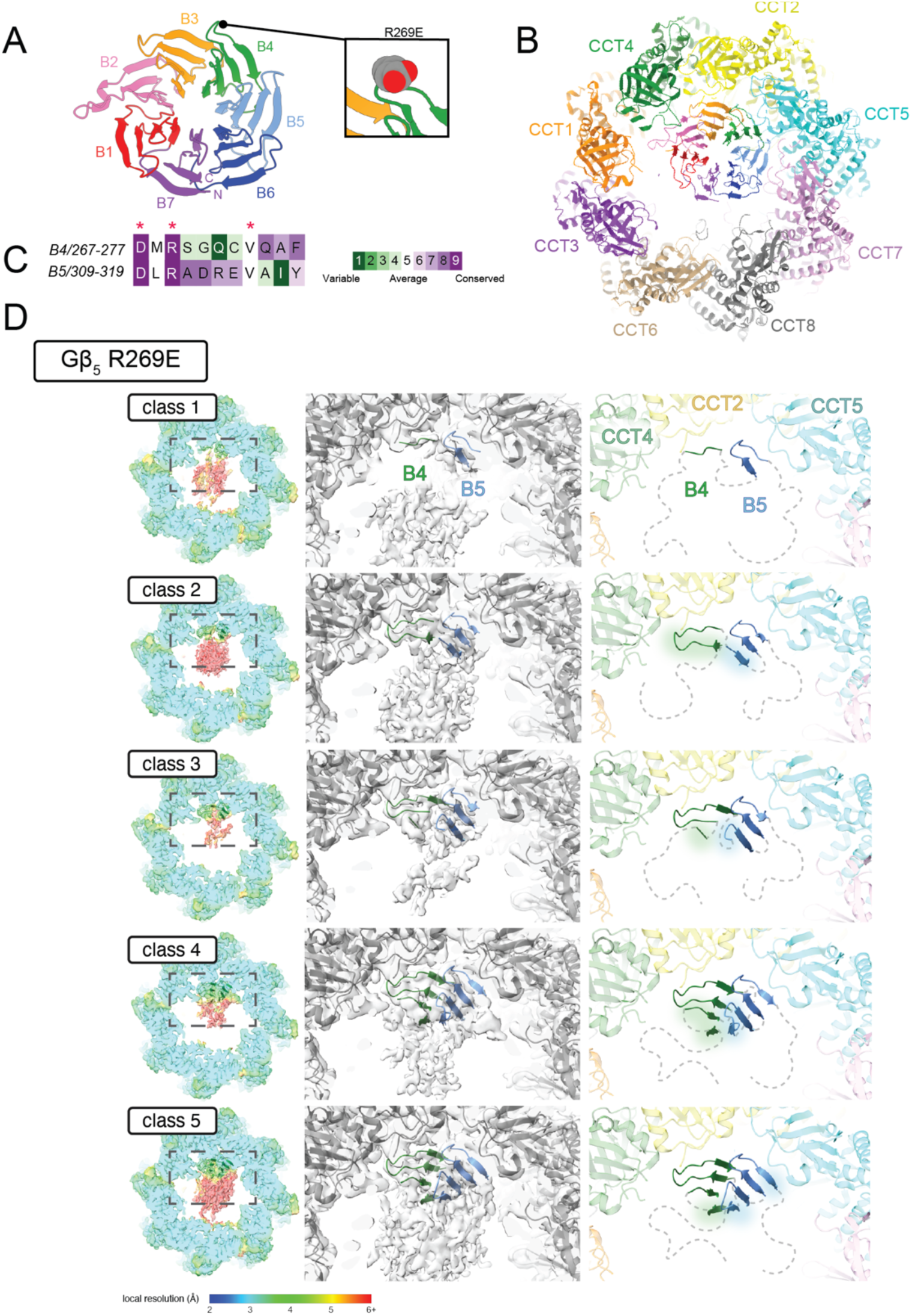
Gβ5 R269E Folding Intermediates. (A) Ribbon structure of Gβ_5_. The same colors for the β-sheet blades are used in all the figures. The inset indicates the location of the R269E mutation. (B) Structure of completely folded Gβ_5_-CCT (PDB 8SGL), showing the contacts of blades 3-5 with CCT2,5, and 7, respectively. (C) Alignment and Consurf conservation analysis of the outer strand of blades 4 and 5. Red asterisks mark residues in blade 4 that contact CCT5 and are conserved in blade 5. (D) Structural analysis of Gβ_5_ R269E reconstructions. Each row depicts an intermediate of Gβ_5_ R269E folding ordered from least to most folded. Left columns show the reconstruction density colored by local resolution with Gβ_5_ R269E structural models in green ribbons. Models were only built into density that exceeded 5 Å local resolution. Lower resolution density was considered unfolded. Middle columns show an inset of the Gβ_5_ R269E models with ribbons colored according to blade number as in Panel A. CCT models are shown as gray ribbons overlayed on the reconstruction density. Right columns show the corresponding inset of structural models. Unstructured regions are indicated as dashed lines. Advances in folding are indicated with a glow effect surrounding the ribbons.

Multiple mutations in Gβ_5_ result in diseases associated with heart arrythmias, neurodevelopmental delays, and retinopathies because of a loss of RGS-Gβ_5_ dimer function (38–47). Among these are several point mutations predicted to disrupt Gβ_5_ folding, including the S123L and the G257E substitutions (Gβ_5_ long splice form amino acid numbering) (38, 40, 44). S123L causes a milder form of disease referred to as Language Delay and ADHD/Cognitive Impairment with or without cardiac arrhythmia (LADCI) (38, 40, 44), whereas G257E results in more severe disease referred to as Intellectual Development Disorder with Cardiac Arrhythmia (IDDCA) (44). Patients with IDDCA present severe phenotypes including intellectual disability, inability to speak, occasionally autism, and arrhythmia that can be fatal. The molecular basis for these differences in disease severity is unknown, but both mutations are found in the core of the β-propeller and would be predicted to disrupt Gβ_5_ folding (40, 44). S123L is located in blade 1, the last blade to fold, while G257E is found in blade 4, the blade that initiates folding (37).

In the current study, we investigated both the importance of electrostatic interactions in initiating Gβ_5_ folding as well as the possible disruption of folding caused by the S123L and G257E mutations. To address folding initiation, we made an R269E substitution that disrupts its electrostatic interaction with CCT5 E256 and determined cryo-EM structures of folding intermediates of this Gβ_5_ mutant bound to CCT. The structures show early folding intermediates attempting to fold by an alternate, off-target pathway from that of wild-type (WT) Gβ_5_, initiating with R311 on blade 5 but stalling early in the process. These findings underscore the importance of the interaction of blade 4 with CCT5 to initiate a productive folding trajectory. To address the effects of the S123L and G257E mutations, we have determined their folding states also by solving cryo-EM structures of folding intermediates. For these mutants, folding initiates normally at blade 4 but stalls partway through the folding process, leaving partially formed β-propellers that are not closed, with S123L achieving a more folded state than G257E. These structures suggest that the mutations cause disease by inhibiting closure of the Gβ_5_ β-propeller and that the folding defect could be overcome by pharmacological chaperones that stabilize the closed form of the β-propeller (48, 49).

## Results

### The R269E substitution disrupts Gβ_5_ folding

Our previous cryo-EM structural analysis of Gβ_5_ folding by CCT identified an electrostatic interaction between Gβ_5_ R269 and CCT5 E256 in the earliest folding intermediate, suggesting that this interaction may initiate Gβ_5_ folding (37). Consistent with this idea, an R269E charge swap substitution increased binding of Gβ_5_ to CCT and dramatically decreased the formation of the Gβ_5_-RGS9 dimer (37), both of which are diagnostic of a folding defect. Additional evidence for a key role of R269 in initiating CCT-mediated Gβ_5_ folding comes from its high evolutionary conservation (Fig. 1C) despite its location on a surface loop not essential to the β-propeller structure. Indeed, the other residues of this loop are much less conserved. Moreover, the strong conservation cannot be explained by R269 contributions to RGS9 binding (50).

To further understand the role of R269 in initiating Gβ_5_ folding, we purified Gβ_5_ R269E bound to CCT and the Gβ co-chaperone PhLP1 directly from HEK-293T cells overexpressing human Gβ_5_ R269E and human PhLP1. We stabilized the folding-competent closed CCT state with ATP and AlF_x_, thereby creating the ATP hydrolysis transition state mimic ADP-AlF_x_, and subjected the complex to single-particle cryo-EM analysis. The reconstruction of the CCT complex in this closed state reached a consensus resolution of 2.7 Å (*SI Appendix* Fig. S1) and revealed density within the central chamber of CCT that prompted further investigation by focused 3D variability analysis (3DVA) and focused 3D classification using cryoSPARC (51). From this classification, five reconstructions ranging between 3.0 to 3.1 Å resolution were determined (*SI Appendix* Fig. S1and S2). All these reconstructions contain density for Gβ_5_ R269E, while only three reconstructions contained partial PhLP1 density at a local resolution better than 5 Å (*SI Appendix* Fig. S1). The PhLP1 density in these reconstructions closely resembles that previously observed for the CCT-PhLP1-Gβ_5_ WT complex (37) and yields no new insights into its role during the substrate folding cycle.

Based on the evidence that R269 initiates Gβ_5_ folding (37), we predicted that the R269E substitution would block folding and leave Gβ_5_ R269E in a molten globule state unable to form the salt bridge needed to trigger folding. However, the five structures from the reconstructions show evidence of partial Gβ_5_ R269E folding that progresses from an initial β-hairpin associated with CCT5 (class 1) to a class that reveals two fully folded blades (class 5) (Fig. 1D). (In this analysis and throughout the study, we considered a region folded if the local resolution exceeded 5 Å.) No completely folded Gβ_5_ β-propellers were observed even though control reconstructions of WT Gβ_5_ using the same procedures with fewer particles produced a completely closed β-propeller (*SI Appendix* Fig. S3A).

### Alternate initiation pathway of Gβ_5_ R269E

The β-hairpin associated with CCT5 in class 1 of Gβ_5_ R269E is reminiscent of the initiating interaction of R269 of blade 4 with E256 of CCT5 in WT Gβ_5_ (Fig. 2A). However, the charge-swapped R269E substitution would disfavor this interaction due to electrostatic repulsion, which prompted us to examine the orientation of Gβ_5_ R269E in our reconstructions more closely. The orientation of blades in the reconstruction was examined by rigid-body fitting of a fully folded Gβ_5_ β-propeller (PDB: 8SGL) (Fig. 1B) into the density of the most folded class (class 5). This model was then rotated such that all combinations of blades were assessed for fit into the density. Blade 5 was the only one with a surface-exposed arginine residue (R311) that fit into the density between Gβ_5_ and CCT5 (Fig. 2A). This orientation places blade 5 into the density where initiation occurs and will be hereafter referred to as the “alternate” orientation to distinguish it from the “native” orientation of WT Gβ_5_ in which blade 4 occupies this position. Models were refined for each class in both the native and alternate orientations, and the final orientation of the model was assigned by visual inspection of its fit into density at a local resolution of 5 Å or better. Pairwise sequence alignment of the outer β-strands of blade 4 and blade 5 reveals similarities between CCT-interacting residues (Fig. 1C). Specifically, D267, R269, and V274 from blade 4 were previously identified as key interactors with CCT5 (37), and these have identical counterparts within blade 5 at D309, R311, and V316, respectively. This alignment, as well as the fit of the alternate model in the two most folded classes, suggests that these residues in blade 5 can be positioned to form the salt bridge, hydrogen bond, and hydrophobic contacts with E256 and W324 of CCT5 that were previously observed with WT Gβ_5_ (Fig. 2A) (37).

**Figure 2.**
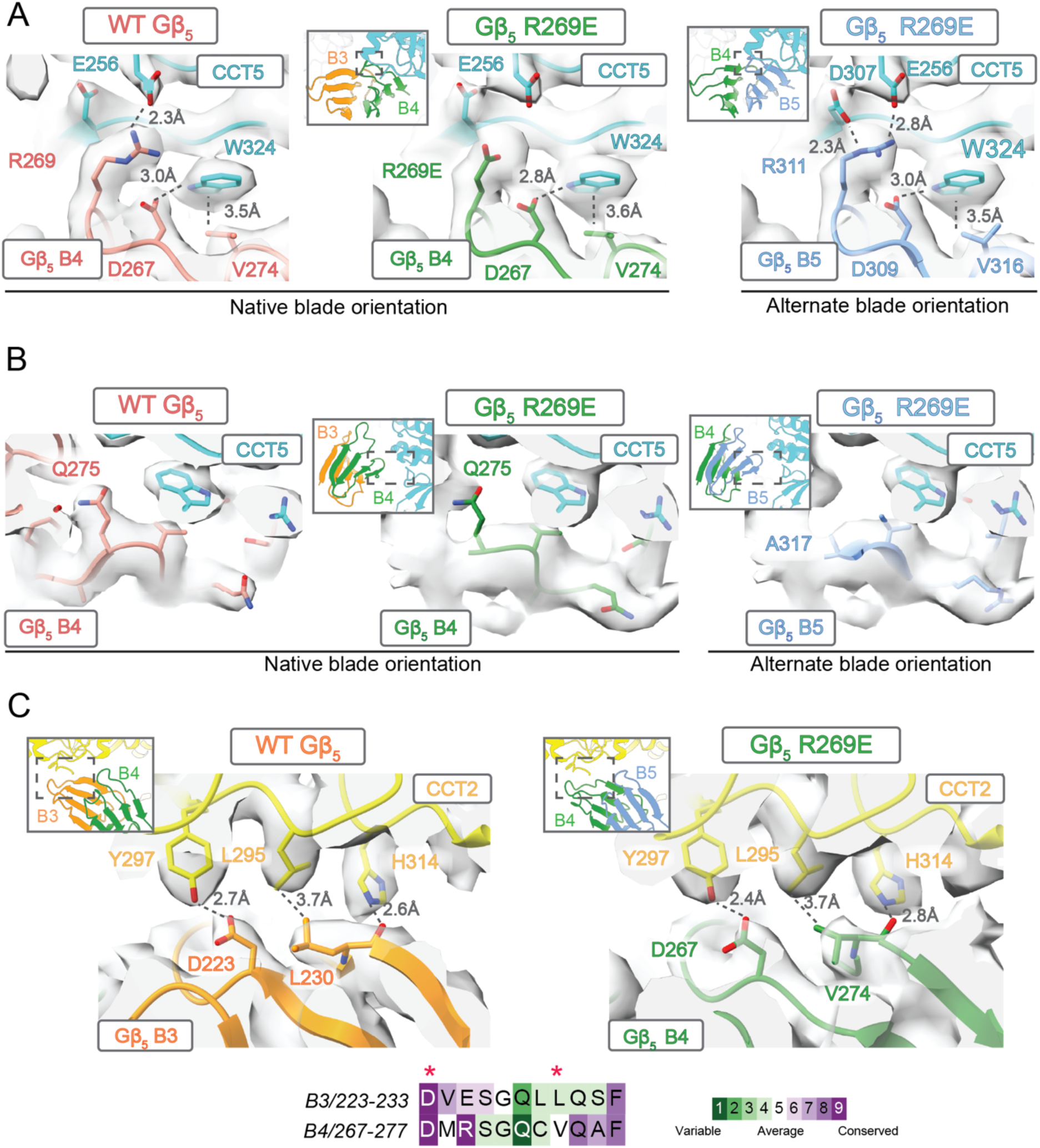
Alternate initiation of Gβ_5_ R269E. (A) Series of electrostatic interactions between CCT5 and blade 4 (B4) of WT Gβ_5_ (left), Gβ_5_ R269E in the native blade orientation (middle), or Gβ_5_ R269E in the alternate blade orientation (right). Insets indicate the location of the region being examined in the complex. (B) Comparison of the fit of Q275 and A317 into the density organized as in (A). (C) Top – interactions between WT Gβ_5_ blade 3 (B3) and CCT2 in the native orientation (left) and between Gβ5 R269E blade 4 and CCT2 in the alternate orientation (right). Insets indicate the location of the region being examined in the complex. Bottom – alignment and Consurf conservation analysis of the outer β-strands of blade 3 and blade 4. Red asterisks mark residues in blade 3 that contact CCT2 and are conserved in blade 4.

Additional comparison of the fits of the native and alternate orientations into the density of Gβ_5_ R269E provides further support for the alternate model. In the WT Gβ_5_ structure, clear bulky density is observed for Q275 of blade 4 in the native orientation. However, the density from the Gβ_5_ R269E reconstruction in this region does not accommodate the glutamine side chain in the native orientation (Fig. 2B), whereas the alternate orientation places a smaller residue A317 of blade 5 in this density that fits well.

Other reconstructions from 3D classification reveal folding intermediates beyond the initial blade. The alternate orientation shifts blade 4 such that it would interact with CCT2 instead of CCT5. We therefore compared the model and density corresponding to the native and alternate interaction between CCT2 with blades 3 and 4, respectively (Fig. 2C). Sequence alignment of the outermost β-strand in blade 3 and blade 4, which constitutes the interaction interface with CCT, reveals several similar residues (Fig. 2C). Notably, both blades contain an aspartate (D223 of blade 3, D267 of blade 4) that can form a hydrogen bond with Y297 of CCT2 (Fig. 2C). Thus, in the alternate orientation, blade 4 can replace blade 3 and make productive contacts with CCT2 that allow blade 4 to fold.

After the formation of two complete blades, folding appears to stall as evidenced by a lack of additional folding intermediates and poor local resolution of surrounding Gβ_5_ density (Fig. 1D). We hypothesize that blades 3 and 6 do not form favorable interactions with CCT4 and CCT7 in the alternate orientation. In WT Gβ_5_, R311 of blade 5 forms a salt bridge with E245 and D297 of CCT7, and several additional hydrogen bonds form between blade 5 and CCT7. However, R311 and much of the outer β-strand of blade 5 is not conserved in blade 6 and rigid body fitting of blade 6 in the alternate orientation show few compensatory interactions, resulting in displacement of blade 6 away from CCT7 (*SI Appendix* Fig. S3B). Moreover, clashes are also predicted to occur between W304 and Q303 of CCT5 and L355 of blade 6 in this orientation (*SI Appendix* Fig. S3C). In the case of blade 3, further inspection of its predicted interaction surface with CCT4 in the alternate orientation reveals potential clashes between Q231 of blade 3 and I304 of CCT4 (*SI Appendix* Fig. S3C). Thus, the alternate orientation appears to stall folding due to the inability of blades 3 and 6 to interact favorably with CCT4 and CCT7, respectively. Together, these data suggest that the complete folding of Gβ_5_ depends on the stabilization of multiple blades by complementary CCT subunits. While aberrant initiation can occur and does induce partial folding, this alternate folding pathway apparently leads to a dead end.

To further test the possibility of alternate Gβ_5_ folding initiation through R311, we prepared a Gβ_5_ R311E mutant and an R269E/R311E double mutant (Fig. 3A) and measured their effects on binding to CCT and RGS9, reasoning that if the alternate orientation were responsible for the residual folding of Gβ_5_, then these mutants might disrupt folding even further. The mutants showed the increased binding to CCT (Fig. 3B) and the dramatically decreased binding to RGS9 (Fig. 3C) observed with the R269E mutant, indicating that the R311E mutants were disrupting Gβ_5_ folding similarly to the R269E mutant, but not more so in these indirect assays. To test folding more directly, we measured ATP-mediated release of the mutants from CCT. The ATPase cycle of CCT drives substrate folding and release, and if a substrate cannot fold, it releases poorly from CCT during this process (52, 53). As expected, ATP addition released 50% of WT Gβ_5_ from CCT in this assay (Fig. 3D). The R269E mutant showed a similar 50% release, while R311E and R269E/R311E showed much less release (∼20%), indicating that the R311E mutations disrupted CCT-mediated Gβ_5_ folding more than R269E.

**Figure 3.**
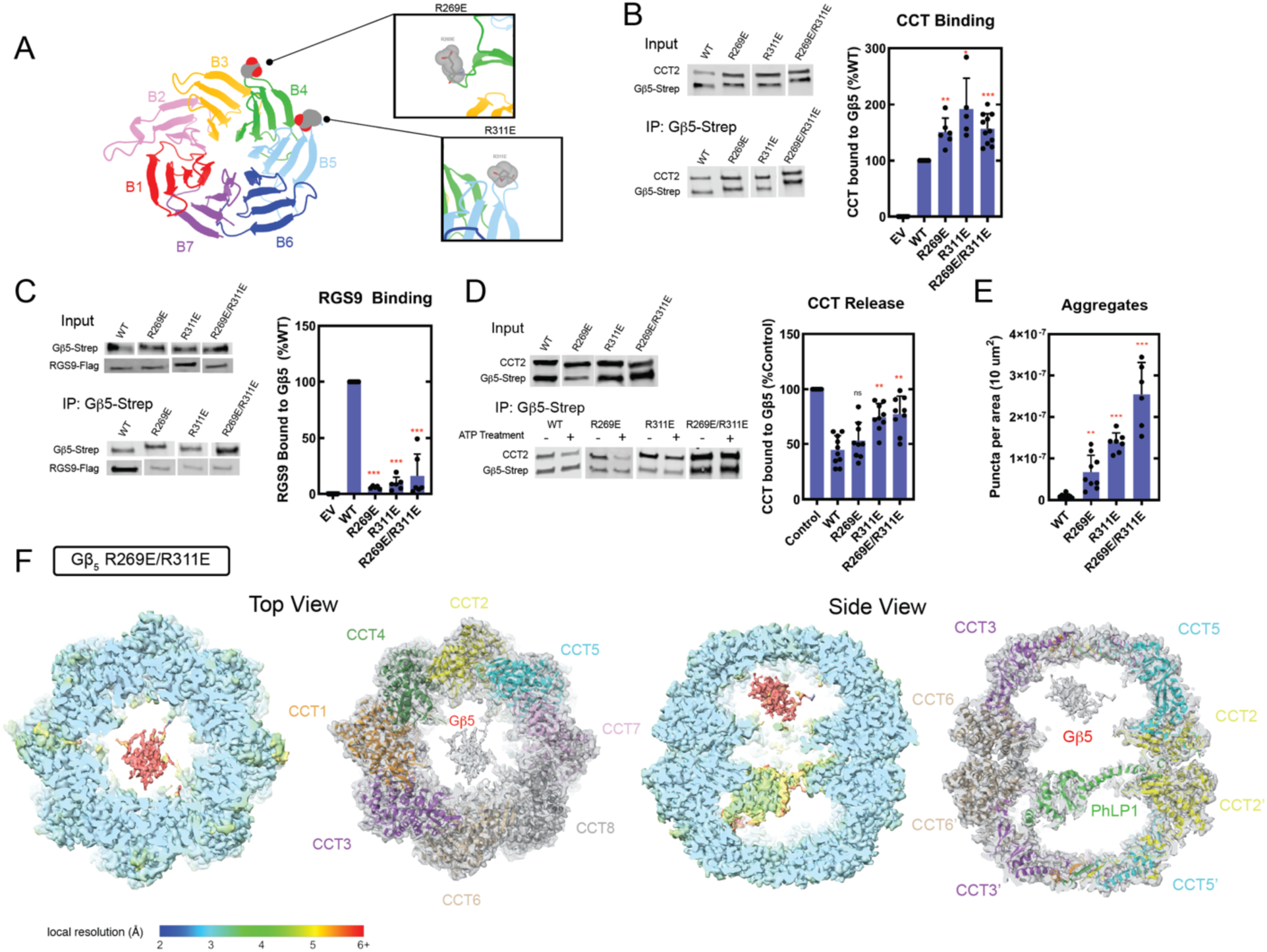
Effects of R269E, R311E and R269E/R311E mutations on Gβ_5_ function. (A) Schematic representation of Gβ_5_ with insets highlighting the R269E and R311E mutations. (B) Binding of CCT to the Gβ_5_ mutants. The CCT/Gβ_5_ ratios in Gβ_5_ immunoprecipitates are shown as a percentage of the WT. (C) Binding of RGS9 to the Gβ_5_ mutants. The RGS9/Gβ_5_ ratios in Gβ_5_ immunoprecipitates are shown as a percentage of the WT. (D) ATP-mediated release of Gβ_5_ from CCT. The CCT/Gβ_5_ ratios after ATP treatment are shown as a percentage of the no ATP control. (E) Aggregation of Gβ_5_ mutants. Quantification of Gβ_5_ puncta in cells over-expressing Gβ_5_ mutants was determined from the immunofluorescence images in *SI Appendix Fig. S4.* Each data point represents a biological replicate indicating the average puncta density from 9 cells of a separately transfected sample. In the all graphs, statistical significance was determined using Welch t-tests compared to WT Gβ_5_ (* p < 0.05, ** p < 0.01, *** p < 0.001, n.s. = not significant). (H) Structure of the CCT-PhLP1-Gβ_5_ R269E/R311E complex. The panel shows top and side views with the reconstruction density colored by local resolution (left) or with the CCT-PhLP1 structural model (PDB 8SH9) docked into the reconstruction density (right).

As a final measure of folding of the Gβ_5_ mutants, we assessed the degree to which they formed aggregates in the cell. Unfolded proteins often form cellular aggregates that appear as puncta in immunofluorescence, so to assess misfolding, we transfected HEK-293T cells with the WT Gβ_5_, R269E, R311E, or R269E/R311E mutants and localized the Gβ_5_ by immunofluorescence labeling. We observed large increases in puncta in the mutants compared to WT, with the most puncta in the double mutant followed by the R311E and then the R269E single mutants (Fig. 3E and *SI Appendix* Fig. S4).

Collectively, these Gβ_5_ folding assays indicate the most disruption of Gβ_5_ folding by the R269E/R311E double mutant followed by the R311E and then the R269E single mutants. These results are consistent with the alternate initiation model, which would be blocked by mutating both R269 and R311. To test this notion further, we directly visualized the folding of the Gβ_5_ double mutant by purifying it in complex with CCT and PhLP1 and solving the structure by cryo-EM. The ATP-AlF_x_ closed structure resolved to an overall resolution of 3.0 Å (*SI Appendix* Fig. S5 and S6) and revealed an unstructured mass attributable to Gβ_5_ in one CCT folding chamber (Fig. 3F). Importantly, no direct contacts of this mass with CCT5 were found nor were any Gβ_5_ folding intermediates observed. This mass closely resembles the unstructured mass prior to folding initiation in the WT Gβ_5_-CCT folding trajectory (37). In contrast, the PhLP1 co-chaperone was clearly visualized in the opposite folding chamber with a local resolution similar to that seen in the WT Gβ_5_-CCT structure (37), indicating the resolution within the folding chambers was optimal. Therefore, the inability of Gβ_5_ R269E/R311E to initiate folding on CCT5 confirms the existence of an alternate folding pathway and demonstrates that by disrupting both the native and alternate folding pathways, Gβ_5_ folding is abrogated.

### Folding of the IDDCA-causing Gβ_5_ G257E mutant

We expanded our examination of Gβ_5_ folding to investigate how pathogenic Gβ_5_ mutations might disrupt the folding process. Among these, we selected two missense mutations that could provide unique insights into Gβ_5_ folding. One was the G257E mutation, which is located on blade 4, the first blade to fold. The other was the S123L mutation located on blade 1, the last blade to fold. G257E introduces a potential folding defect by inserting a negative charge into the center of the β-propeller at the interface between blades 4 and 5 (Fig. 4A) and causes the more severe form of Gβ_5_ deficiency disease (IDDCA). In contrast, S123L adds a bulky side chain at the interface between blades 1 and 2 of the β-propeller and causes the less severe Gβ_5_ deficiency disease (LADCI). G257 and S123 are well conserved among Gβ_5_ proteins (Fig. 4B), which is consistent with the tight packing of these residues in the core of the β-propeller (Fig. 4A) but cannot be attributable to interactions with RGS9 because these residues do not contact RGS9.

**Figure 4.**
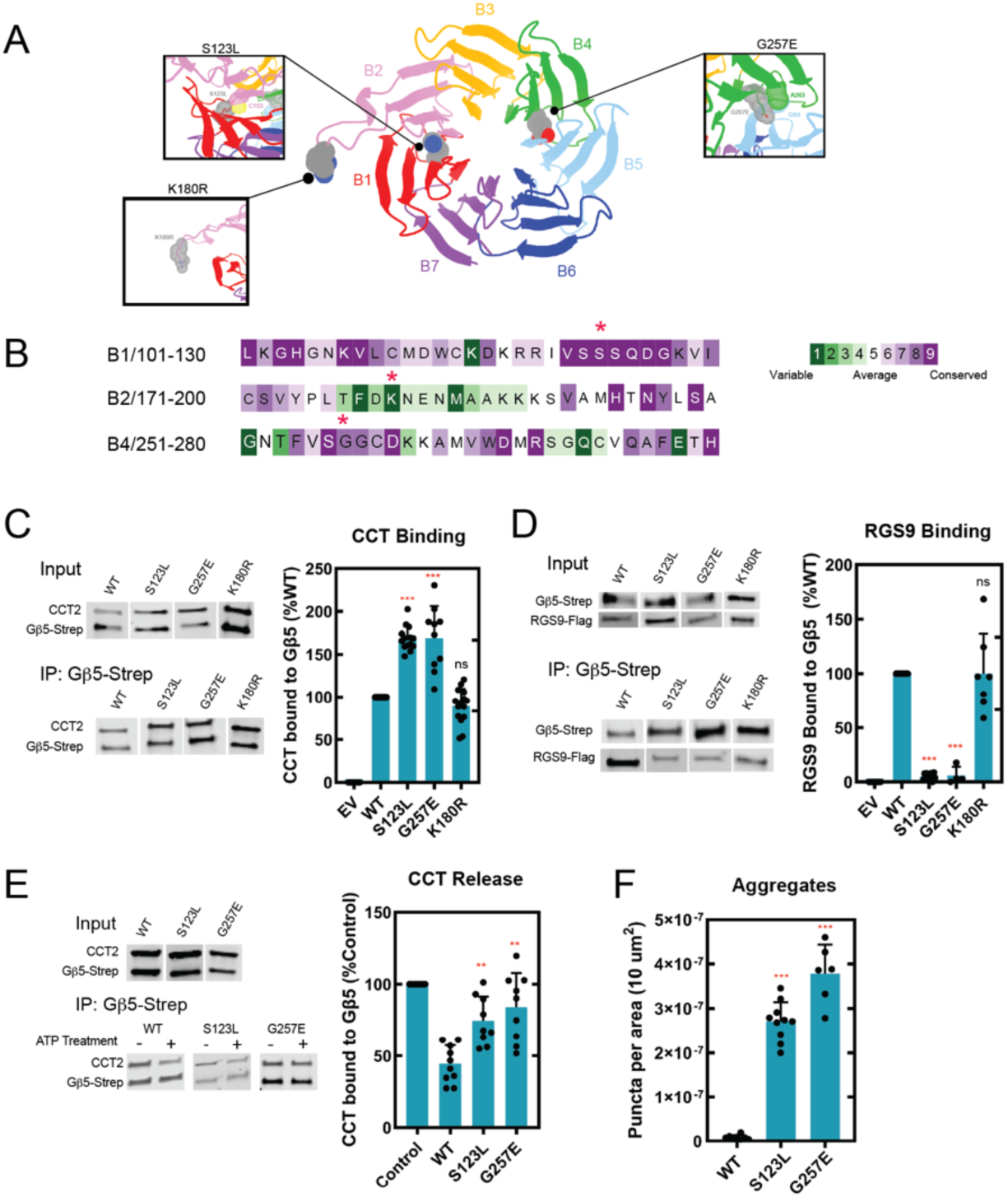
Effects of disease-causing mutations on Gβ_5_ function. (A) Schematic representation of Gβ_5_ with insets highlighting the S123L and G257E mutations as well as the K180R variant and their predicted clashes with neighboring blades. (B) Consurf conservation analysis of the β-propeller blades containing S123, K180, and G257 (marked by asterisks). (C) Binding of CCT to the Gβ_5_ mutants. The CCT/Gβ_5_ ratios in Gβ_5_ immunoprecipitates are shown as a percentage of the WT. (D) Binding of RGS9 to the Gβ_5_ mutants. The RGS9/Gβ_5_ ratios in Gβ_5_ immunoprecipitates are shown as a percentage of the WT. (E) ATP-mediated release of Gβ_5_ from CCT. The CCT/Gβ_5_ ratios after ATP treatment are shown as a percentage of the no ATP control. (F) Aggregation of Gβ_5_ mutants. Quantification of Gβ_5_ puncta in cells over-expressing Gβ_5_ mutants was determined from the immunofluorescence images in *SI Appendix Fig. S7.* In the all graphs, statistical significance was determined using Welch t-tests compared to WT Gβ_5_ (* p < 0.05, ** p < 0.01, *** p < 0.001, n.s. = not significant).

To test the effects of these mutants on Gβ_5_ function, we first measured their binding to CCT and to RGS9 and found that both exhibited functional defects consistent with disrupted folding like the R269E mutant (37). Both S123L and G257E displayed increased binding to CCT and markedly decreased binding to RGS9 compared to WT (Fig. 4C-D). As a negative control, we tested binding of another Gβ_5_ variant with a conservative substitution (K180R) in a variable loop that is not associated with disease. The K180R variant showed binding to CCT and RGS9 that was very similar to the WT (Fig. 4C-D). This result rules out artifacts from the mutagenesis process that could disturb binding.

In the ATP release assay (Fig. 4E), both S123L and G257E released considerably less Gβ_5_ from CCT (20%) than WT (50%), adding further support for a folding defect in these mutants. Finally, immunofluorescence imaging of cells transfected with the mutants showed many more puncta than cells transfected with WT Gβ_5_ (Fig. 4F and *SI Appendix* Fig. S7), pointing to disrupted folding leading to aggregation in the mutants. All these findings indicate that the S123L and G257E mutations cause folding defects in Gβ_5_.

To understand these folding defects at the molecular level, we purified the CCT-PhLP1-Gβ_5_ G257E and the CCT-PhLP1-Gβ_5_ S123L complexes and determined cryo-EM structures of their folding intermediates. For G257E, the consensus structure of closed state CCT reached a resolution of 2.9 Å and contained Gβ_5_ density within the central chamber (*SI Appendix* Fig. S8). Focused 3DVA and 3D classification over the Gβ_5_ density was performed and yielded six maps with resolutions ranging between 3.2-3.3 Å (*SI Appendix* Figs. S8-S9). Within these maps, the β-propeller blade orientation was assigned by rigid-body fitting as described above for the R269E complex. The resulting structures were ordered from least to most folded, revealing a folding pattern that initiates like WT Gβ_5_ but does not progress beyond a partially folded β-propeller (Fig. 5). As with WT Gβ_5_, folding begins with the formation of a β-hairpin of blade 4 (class 1) followed by formation of a β-hairpin of blade 3 (class 2). Folding continues until three β-strands of each blade 3 and blade 4 are formed (class 3). These classes comprise 45% of all closed state particles. The remaining 55% of particles show fully folded blades 3 and 4 with incremental folding of blades 2 and 5 until, in the final class, two β-strands of blade 2 and a β-hairpin of blade 5 are folded (Fig. 5). As with R269E, this partially folded structure was not a result of a limited particle number as a similar analysis of WT Gβ_5_ with fewer particles produced a completely closed β-propeller (*SI Appendix* Fig. S3A).

**Figure 5.**
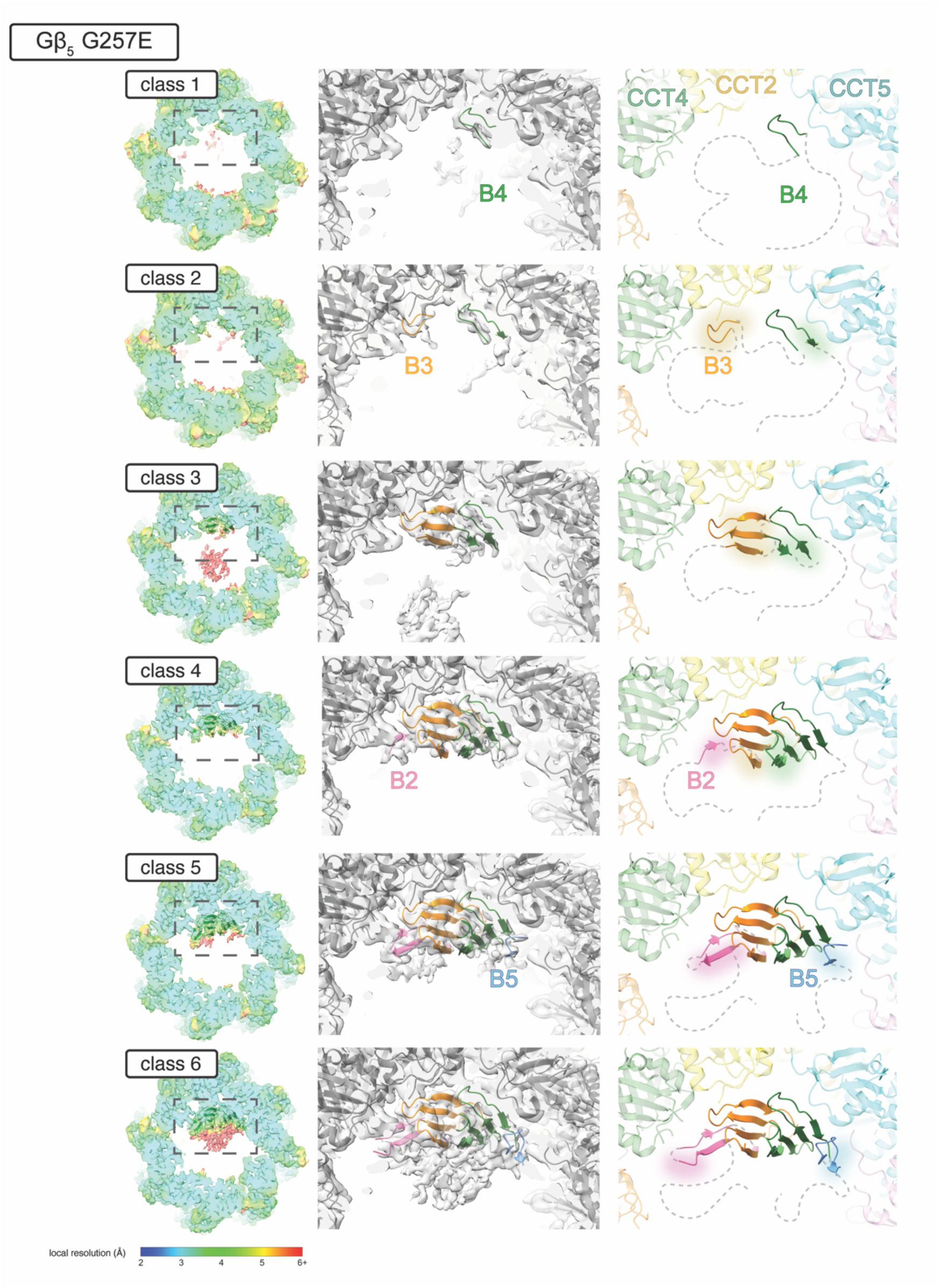
Gβ_5_ G257E Folding Intermediates. Each row depicts an intermediate in Gβ_5_ G257E folding ordered from least to most folded. Views in the three columns are as in Figure 1.

The G257E mutation is located within the second inner β-strand of blade 4, and the side chain is exposed at the interface between blades 4 and 5 (Fig. 4A). Accordingly, we observe that folding initiation progresses similarly to WT Gβ_5_ but appears to stall with the folding of blade 5, leaving most of the particles in classes where blade 5 begins but does not complete folding (Fig. 5). This disruption of folding is consistent with the steric hindrance expected by the replacement of a glycine at the blade 4-blade 5 interface with the larger glutamate residue.

### Folding of the LADCI-causing Gβ_5_ S123L mutant

The folding intermediates of Gβ_5_ G257E show how a pathogenic mutant can hinder folding at early stages but leave open the question of how mutations in blades that fold later might disrupt the advanced stages of folding. To address this question, we determined the structure of the CCT-PhLP1-Gβ_5_ S123L complex via cryo-EM. A consensus structure for the closed state was solved to a resolution of 2.9 Å (*SI Appendix* Fig. S10), and focused 3DVA and 3D classification yielded five reconstructions with resolutions ranging between 3.1-3.2 Å (*SI Appendix* Figs. S10 and S11). These structures show that Gβ_5_ S123L folding also initiates at blade 4, progresses through blade 3 and two strands of blade 2 on one side of the β-propeller, and continues along the other side until blade 5 is completely folded (Fig. 6), resulting in an approximately half-folded β-propeller. These folding intermediates are similar to those of Gβ_5_ G257E, except that in the most folded G257E intermediate, blade 5 folds minimally. Again, partial folding was not caused by limited particle numbers in the S123L dataset because WT Gβ_5_ with fewer particles produced a fully closed β-propeller (*SI Appendix* Fig. S3A). Collectively, these findings demonstrate that the S123L mutation not only blocks folding of the blade in which it is found (blade 1), but it also destabilizes the folding of the other β-propeller blades that do not contact CCT (blades 2, 6, and 7).

**Figure 6.**
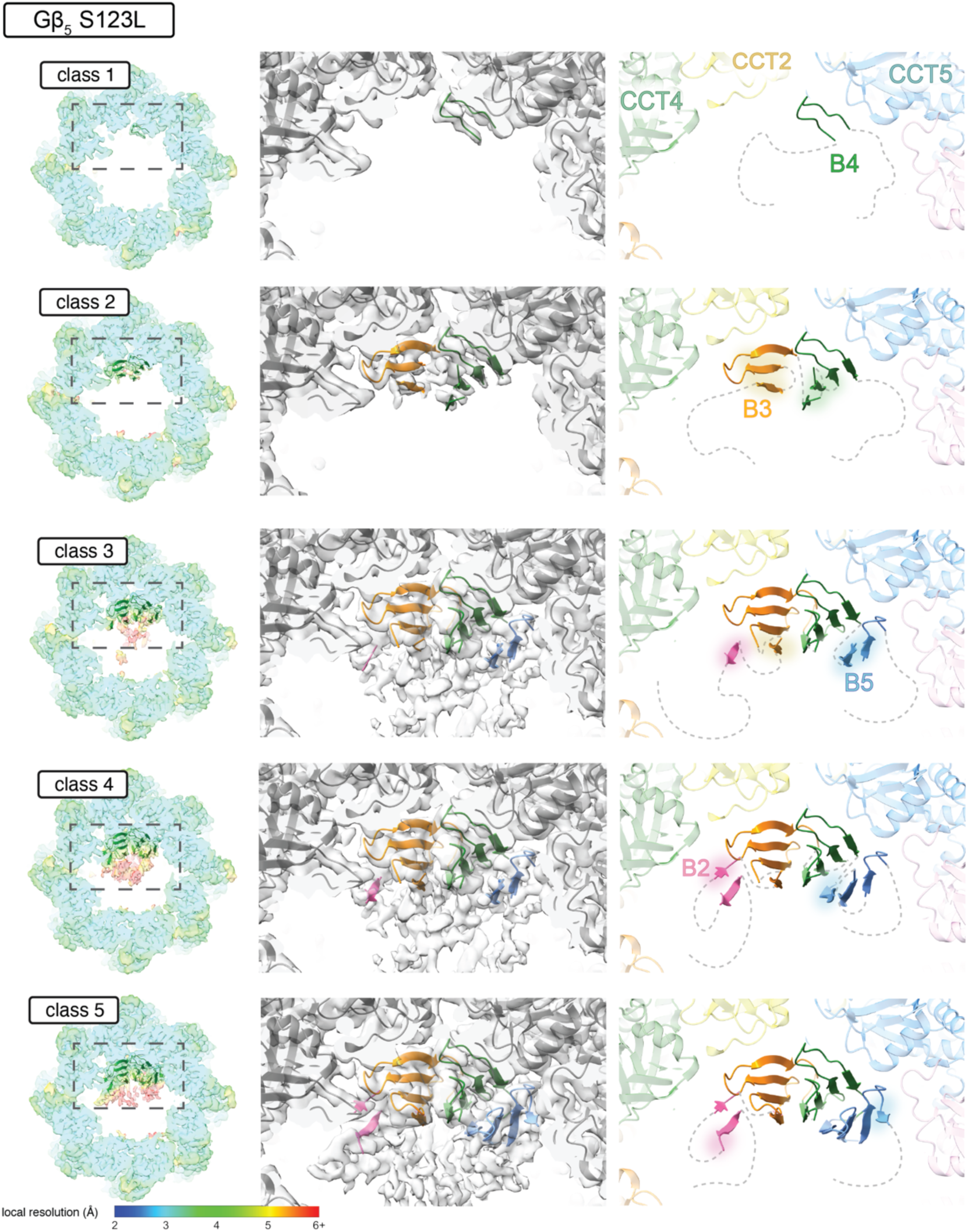
Gβ_5_ S123L Folding Intermediates. Each row depicts an intermediate in Gβ_5_ S123L folding ordered from least to most folded. Views in the three columns are as in Figure 1.

## Discussion

Our previous work to visualize the CCT-mediated folding trajectory of WT Gβ_5_ by cryo-EM provided structural and biochemical evidence that the electrostatic interaction between Gβ_5_ R269 and CCT5 E256 plays a key role in initiating Gβ_5_ folding (37). This finding raised questions about why R269 in Gβ_5_ blade 4 was the most favorable initiation site even though other arginine residues on the surface of Gβ_5_ could potentially perform the same function. The biochemical and structural findings presented here effectively answer this question. An alternative initiation site exists at R311 of Gβ_5_ blade 5. This site harbors the same key CCT5 contact residues as those of blade 4 (Fig. 1B), which allows blade 5 to also initiate folding on CCT5. However, while initiation at blade 5 moves blade 4 into position to interact favorably with CCT2 (Fig. 2C), it also places blades 3 and 6 next to CCT4 and CCT7, respectively, with few favorable interactions and some significant steric clashes (*SI Appendix* Figs. 3B and 3C). Consequently, the Gβ_5_ β-propeller becomes misaligned with the CCT subunits needed for folding, and blades 3 and 6 do not fold. These observations suggest that WT Gβ_5_ might toggle between initiating interactions of blade 4 or blade 5 with CCT5, but subsequent productive interactions with the other CCT subunits favors initiation at blade 4. Thus, specificity of Gβ_5_ folding by CCT appears to derive from multiple interactions of Gβ_5_ blades 3, 4, and 5 with CCT subunits 2, 5, and 7, respectively, which highlights the importance of the multifaceted interaction landscape within the interior of the CCT folding chamber to support protein folding.

In addition to alternate initiation, R311 may have other important functions in Gβ_5_ folding as evidenced by the severe disruption caused by the R311E mutation alone (Fig. 3). This function is likely to stabilize blade 5 via electrostatic interactions of R311 with E245 and D297 of CCT7 (37) and allow folding to propagate through blade 5. Notably, R311 does not initiate folding through these interactions with CCT7 because none of the structures show initiation on CCT7, only on CCT5. Thus, R311 make essential contributions to the folding of the Gβ_5_ β-propeller by providing an alternate initiation site on CCT5 and by assisting in propagating folding through blade 5 on CCT7. Remarkably, the R269E and the R311E mutations, which are located on surface loops not involved in the formation of the β-propeller core, are enough to completely disrupt Gβ_5_ folding. This finding attests to the importance of CCT in forming the Gβ_5_ β-propeller.

Missense mutations that cause single amino acid changes are responsible for 30% of all genetic diseases (54), and understanding the molecular defects caused by these mutations can lead to treatments for the underlying disfunctions (55, 56). A recent study estimates that 60% of missense mutations disrupt protein stability (57), raising questions about how the cellular proteostasis network deals with mutations that disrupt protein folding. These questions motivated our structural analysis of the Gβ_5_ G257E and S123L mutations to determine if these amino acid substitutions cause misfolding of the Gβ_5_ β-propeller. The G257E mutation occurs on blade 4 at the interface between blades 4 and 5 (Fig. 4A). The structures of the G257E folding intermediates (Fig. 5) show that the mutant does not interfere with initiation on blade 4 nor the folding of blade 3, but folding does not progress through blade 5. The inability of blade 5 to form indicates that the mutation destabilizes the inter-blade contacts between blades 4 and 5. These contacts appear necessary for blade 5 to fold, even though the stabilizing interactions of blade 5 with CCT7 remain intact. This observation argues that CCT orchestrates Gβ_5_ folding by stabilizing folding intermediates, but these interactions are insufficient to complete folding if the core of the protein is unable to achieve a stable conformation.

In the case of S123L, the mutation is found in blade 1 at the interface between blades 1 and 2 (Fig. 4A). These blades are the last to form as the β-propeller closes in the Gβ_5_ folding trajectory (37). The lack of folding of blades 1 and 2 is consistent with disruption of the interface between these blades, which inhibits the β-propeller from closing. Notably, the mutation not only destabilizes blades 1 and 2 but also blades 6 and 7, which are the four blades that do not interact directly with CCT. This observation indicates that these blades derive their stability from interactions with each other as the β-propeller closes and that an inability to close destabilizes all four blades.

The G257E mutation causes a significantly more severe form of Gβ_5_ deficiency disease (IDDCA) than the S123L mutation (LADCI), and the disruption of folding caused by G257E is greater than S123L. With G257E, only blades 3 and 4 are fully folded while with S123L all three blades that contact CCT (blades 3-5) are clearly folded (Figs. 5 and 6). However, neither of these Gβ_5_ mutants bound RGS9 effectively, they each resisted ATP-mediated release from CCT similarly, and both formed cellular aggregates to a similar extent. Thus, these data do not offer a compelling explanation why disease severity is greater with G257E. Perhaps the G257E mutation induces a deleterious gain of function not observed here.

In conclusion, these structural studies support the validity of the WT Gβ_5_ folding trajectory reported previously (37) and identify the molecular defects in folding from Gβ_5_ mutants as illustrated in Fig. 7. The inability of the R269E mutant to initiate the native folding pathway and the specific disruptions caused by the S123L and G257E mutations support the previous observation that folding initiates on blade 4 and progresses to adjacent blades in both directions around the β-propeller. The stalled folding trajectories of these mutants suggest potential therapeutic strategies to stabilize the β-propeller fold using pharmacological chaperones. Therapeutically stabilized mutants could fold more efficiently, allowing more stable Gβ_5_-RGS9 dimers to assemble and perform their Gα-GTP hydrolysis function. A recent extensive study to pharmacologically target β-propeller proteins provides proof of principle that these β-propellers are viable therapeutic targets (48, 49). Finally, while these observations provide a framework for predicting how other missense mutants of Gβ_5_, such as L101P, S123W, and R288Q, may have altered folding, future studies will be needed to determine how misfolding mutations alter folding in other CCT substrates, including actin or tubulin. For example, multiple missense mutations in tubulin are predicted to cause folding defects (58) that could also be revealed by examining the folding intermediates of these mutants.

**Figure 7.**
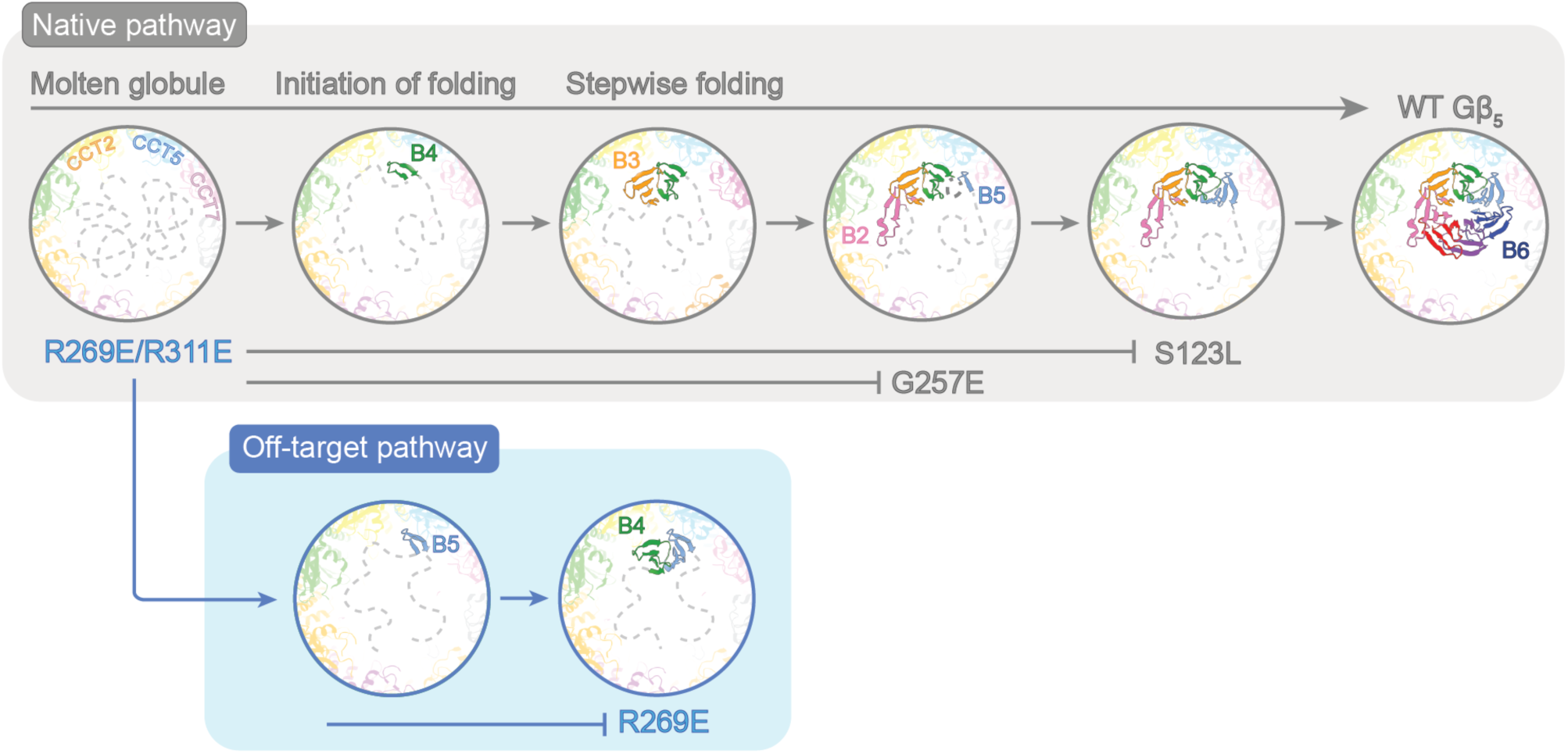
Model of Gβ_5_ mutant folding. The model depicting key steps in the folding of Gβ_5_, including the inability of the R269E/R311E double mutant to initiate folding, the off-target pathway for the R269E mutant, and the point where folding is disrupted by the S123L and G257E mutants.

## Materials and Methods

### Cell culture and transfections

HEK-293T cells were cultured in DMEM/F-12 supplemented with 10% FBS in T-25 flasks. At 40-50% confluency, cells were transfected with 560 µl of a mixture of 510 μl OptiMEM, 30 μl Polyethyleneimine (PEI), and 20 μl of cDNA containing 2 µg of an N-terminal 2x strep peptide-tagged human WT Gβ_5_, Gβ_5_ R269E, Gβ_5_ R311E, Gβ_5_ R269E/R311E, Gβ_5_ G257E, or Gβ_5_ S123L cDNA construct as indicated, 2 µg of a C-terminal myc-His_6_ tagged human PhLP1, 2 µg of a C-terminal Flag tagged human RGS9 cDNAs and 1 μg of GFP cDNA (as a transfection control) all in the pcDNA3.1 vector.

### Immunoprecipitation and immunoblotting

Cells transfected with Gβ_5_ mutants and RGS9 in T-25 flasks were washed in phosphate-buffered saline (PBS), then harvested and lysed in 500 µl of lysis buffer (PBS supplemented with 1% IGEPAL, 0.5 mM PMSF and Halt Protease Inhibitor Cocktail). Lysates were triturated 10 times through a 25-gauge 7/8” needle, then centrifuged at 21,000 x g for 10 minutes at 4°C to clarify the lysates. The lysate protein concentration was measured by BCA assay. Gβ_5_ was immunoprecipitated from the lysate by adding 400 μg of cell lysate to 25 µl of Strep-Tactin resin and incubating for 1 hour at 4°C with inversion. Immunoprecipitates were washed three times in lysis buffer, then resuspended in 40 µl of SDS-PAGE sample loading buffer. Of this, 8 μl samples were loaded and separated on SDS-PAGE on 4-20% gradient gels and transferred to nitrocellulose. The nitrocellulose was probed for 16 hours at 4°C with the indicated primary antibodies at the following dilutions in TBS blocking buffer (LI-COR): Strep (Genscript, 1: 5,000 dilution), c-myc (Invitrogen, 1: 1,000), CCT2 (Abcam, 1: 10,000), and Flag (Sigma Aldrich, 1:1,000). Blots were washed 3 times with TBS-Tween wash buffer (50 mM Tris•HCl pH 7.6, 150 mM NaCl, 50 mM NaOH, 0.05% Tween 20) and probed with appropriate IRDye-labeled secondary antibodies at 1:10,000 dilutions in TBS-Tween wash buffer: IRDye 680RD Goat Anti-Mouse (LI-COR) and IRDye 800RD Goat Anti-Rabbit (LI-COR). Blots were imaged using a LI-COR Odyssey infrared scanner, and band intensities were quantified with the LI-COR Image Studio software. The co-immunoprecipitated CCT or RGS9 bands were normalized to the Gβ_5_ band in each immunoprecipitation. These ratios were then compared as a percentage of the WT ratio. Statistical significance was determined using unpaired t-tests with the Welch correction for unequal variance in GraphPad Prism.

### ATP release

To measure the effects of ATP on the interaction of Gβ_5_ with CCT, we transfected cells with 2x Strep tagged Gβ_5_, myc-His_6_ tagged PhLP1, and Flag tagged RGS9. The lysate was then divided in two sets of 400 μg of protein and immunoprecipitated as described above. The immunoprecipitate was washed twice with lysis buffer and then incubated in lysis buffer containing 10 mM KCl, 5 mM MgCl_2_, ± 5 mM ATP at 4°C with inversion for 30 minutes. The immunoprecipitate was washed twice more and then eluted in 40 μL of SDS-PAGE sample loading buffer. Of this, 8 μl samples were loaded and separated by SDS-PAGE on 4-20% gradient gels and transferred to nitrocellulose. The blots were probed and imaged as described above. The co-immunoprecipitated CCT bands were normalized to the Gβ_5_ band in each immunoprecipitation. These ratios were then compared as a percentage of the no ATP ratio. Statistical significance was determined using unpaired t-tests with the Welch correction for unequal variance in GraphPad Prism.

### Immunolocalization

HEK**-**293T cells were seeded onto coverslips previously treated with 0.01% poly-lysine in 12-well plates and transfected with Gβ5 and RGS9 using 62 μL of the same transfection mixture described above. After 48 hours, cells were washed 3 times with PBS at 37°C, fixed with 4% paraformaldehyde (w/v) in PBS for 15 minutes, and washed 3 times with PBS at 23°C. Cells were then permeabilized with 0.5% Tween 20 (v/v) in PBS for 15 minutes, washed as above, and blocked with 0.05% Goat Serum (v/v) in permeabilization buffer for 15 minutes. Primary antibodies were diluted in blocking buffer, added to the cells, and incubated overnight at 4°C. Strep-Gβ_5_ was detected with anti-Strep mouse antibody (Genscript, 1:250 dilution), and CCT was detected with an anti-CCT2 rabbit antibody (Abcam, 1:500 dilution). Cells were then washed twice with PBS and incubated in blocking buffer for 15 minutes. Secondary antibodies were diluted 1:250 in 1 mg/mL BSA in permeabilization buffer. For the Gβ_5_ puncta quantification, Alexa Fluor 594 anti-mouse (Abcam) was used as secondary antibody. After incubation at room temperature for 1 hour in the dark, the cells were washed 3 times with PBS and mounted with DAPI mounting media (Abcam). After incubation coverslips were mounted using ProLong Gold Antifade (ThermoFisher) as mounting media. Images were acquired using a Leica TCS SP8 confocal microscope using 405 and 561 nm lasers with 60X oil immersion objective lens.

### Image quantification

For puncta analysis, images were processed using Fiji software (59) with a macro designed for puncta quantification (60). The red (Gβ_5_) and blue (DAPI) channels were split in Fiji to yield a binary format from which the background was subtracted. The puncta density from the Gβ_5_ binary images were then quantified in a semi-automated manner in Fiji. The macro tool quantified the average number of puncta per 10 μm^2^ from three cells per image and three images per sample at a prominence value of ‘Find maxima’ set to 50. Each data point in Fig. 3E represents this average from a separately transfected biological replicate. The results were plotted and compared to WT for statistical significance using an unpaired t-test with Welch correction for unequal variance in GraphPad Prism.

### Purification of CCT-PhLP1-Gβ_5_ mutants

For the R269E, R269E/R311E, and G257E mutants, Expi-293F cells were cultured in suspension in 50 mL Expi293 Expression Medium in 250 mL plain-bottomed flasks with shaking at 37°C and 8% CO_2_. At a density of 3.0 × 10^6^ cells per mL and over 95% viability, cells were transfected with a 6 mL mixture containing the ExpiFectamine 293 transfection reagent, 12 µg of a C-terminal myc-His_6_ tagged human WT PhLP1 in the pcDNA3.1 vector, and 12 µg of the N-terminal 2x Strep peptide-tagged human Gβ_5_ mutant constructs in the pcDNA3.1 vector. Cells were fed 18 hours after transfection with the enhancers required for the transfection reagent and harvested 24 hours after feeding.

For the S123L mutation, HEK-293T cells were cultured in DMEM/F-12 supplement with 10% FBS in T-25 flasks. At 40% confluency, cells were transduced with pLenti-Puro virus containing human Gβ_5_ S123L with N-terminal 2X Strep and HPC4 tags. The cells were then treated with puromycin for one week to select for transduced cells. The selected cells were then transduced with pLenti-Blast virus containing human PhLP1 with N-terminal His_6_ and myc tags. The cells were then treated with blasticidin for another week to select for transduced cells. The selected cell line was frozen in aliquots and used as a stable cell line for CCT-PhLP1-Gβ_5_ S123L purification. HEK-293T cells stably expressing Strep-Gβ_5_ S123L and His-PhLP1 were grown to 80% confluency in T-175 tissue culture flasks.

Cells for all conditions were then harvested and lysed in lysis buffer at 5 mL of buffer per gram of harvested cell pellet. For each Gβ_5_ mutant, the lysate was centrifuged at 30,000 x g for 20 minutes and filtered through a 0.45 μm filter followed by a 0.22 μm filter. The CCT-PhLP1-Gβ_5_ complexes were then purified at 4°C using a tandem affinity approach. The filtered lysate was passed for one hour over a HisTrap HP 5 mL column equilibrated with 20 mM HEPES pH 7.5, 50 mM NaCl, 25 mM imidazole, 0.05% CHAPS, and 1 mM TCEP. The column was then washed with 5 column volumes of equilibration buffer, and protein was eluted with linear gradient of 8 column volumes of 25 mM to 500 mM imidazole in equilibration buffer. Eluted fractions were analyzed by SDS-PAGE and Coomassie staining. Fractions containing PhLP1 and CCT were combined and loaded onto 3 mL of Strep-Tactin resin equilibrated with 20 mM HEPES pH 7.5, 20 mM NaCl for an hour. The column was washed three times with 1 column volume per wash of 20 mM HEPES pH 7.5, 50 mM NaCl, and then three times with 1 column volume per wash of 20 mM HEPES pH 7.5, 20 mM NaCl. The CCT-PhLP1-Gβ_5_ complexes were eluted with 3 column volumes of 20 mM HEPES pH 7.5, 20 mM NaCl, 10 mM D-desthiobiotin, 1 mM TCEP, and 5 mM MgCl_2_ and concentrated to 1.5-2.0 μg/μL using a 30 kDa cutoff filter and analyzed by SDS-PAGE and Coomassie staining and immunoblotting. A more detailed protocol has previously been described for the purification, sample preparation, imaging, and reconstruction of the WT CCT-PhLP1-Gβ_5_ complex (61).

### Inducing the closed conformation of CCT

To 10 µl of purified CCT-PhLP1-Gβ_5_ at 1.5-2.0 mg/mL, 0.5 μL of 600 mM KCl, and 1.5 µl each of 10 mM Al(NO_3_)_3_, 60 mM NaF, and 10 mM ATP were sequentially added in rapid succession to avoid Al(OH)_3_ precipitation. The complex was then incubated at 37°C for an hour and centrifuged at 21,000x g for 1 min to remove any precipitate.

### Cryo-electron microscopy specimen preparation

UltrAuFoil R2/2 200 grids (SPT Labtech) were glow discharged with a PELCO easiGlow unit (Ted Pella, Inc.) for 1 minute on each side at 25 mA. 3.5 μL of the closed CCT-PhLP1-Gβ_5_ complex was applied to the glow discharged grid and blotted for 1.5-2 s using a Vitrobot Mk IV (Thermo Fisher Scientific). The grid was plunge frozen into liquid ethane and stored in liquid nitrogen until cryo-EM data collection. Data for the R269E, G257E, and S123L mutants were collected at the University of Utah Electron Microscopy facility on a 300kV Titan Krios G3 with a K3 direct detector (Thermo Fisher Scientific) and the BioQuantum Energy Filter (Gatan) using SerialEM. The nominal magnification was 81,000x corresponding to a pixel size of 1.058 Å. For the CCT-PhLP1-Gβ_5_ R269E complex, three datasets were collected at a dose range between 33 to 43 e/Å^2^ and 40 frames per movie yielding a total of 15,158 micrographs. Three datasets were also collected for the CCT-PhLP1-Gβ_5_ G257E complex using a dose range from 42 to 43 e/Å^2^and 40 frames per movie which resulted in a total dataset size of 22,460 micrographs. Four datasets were collected for the CCT-PhLP1-Gβ_5_ S123L complex using a dose range between 43.01 to 45.41 e/Å^2^ and 40 frames per movie resulting in a total of 21,754 micrographs.

For the R269E/R311E mutant, data were collected at the Brigham Young University Electron Microscopy facility on a 300kV Titan Krios G4 with a Falcon 4 Direct Detector (Thermo Fischer Scientific) and the Selectris X Energy Filter (Thermo Fischer Scientific) using EPU. The nominal magnification was 130,000x corresponding to a pixel size of 0.95Å. Three datasets were collected at a dose range of 25 to 34 e/Å^2^ which resulted in a total dataset size of 37,271 micrographs in the .eer format.

### Image processing

All image processing was performed in cryoSPARC v4.6.2 (51), using a workflow described previously (61). Briefly, for the CCT-PhLP1-Gβ_5_ R269E complex, 15,158 movies were motion corrected, CTF estimated, and curated with a 5Å CTF cutoff. 4,167,017 particles were selected with a blob picker using a radius of 180-220Å. Particles were then extracted with a box size of 300 pixels, Fourier cropped to a box size of 150 pixels, and iteratively 2D classified until 1,168,931 particles remained. These particles were used for Ab initio reconstruction and heterogenous refinement. 643,270 were sorted into the open state and 137,763 particles were junk particles. The remaining 386,973 particles were sorted into the closed state, extracted to the full 300-pixel box size, and then subject to non-uniform refinement. From this consensus structure, 3DVA was conducted with a focused mask over the substrate folding chamber. The component showing the greatest degree of continuous motion was selected for further focused 3D classification and non-uniform refinement, using a mask over the half of the CCT folding chamber containing Gβ_5_. This workflow resulted in six classes with resolutions between 3.0 to 3.1 Å. Five of these classes were distinct Gβ_5_ folding intermediates and were ordered based on the completeness of Gβ_5_ folding.

The image processing for the CCT-PhLP1-Gβ_5_ R269E/R311E, G257E and S123L complexes followed a similar workflow. The final number of closed particles for the CCT-PhLP1-Gβ_5_ G257E complex was 453,385. These particles were further classified via focused 3DVA and focused 3D classification, as above, into six distinct Gβ_5_ folding intermediates with resolutions ranging from 3.1 to 3.2 Å. The CCT-PhLP1-Gβ_5_ S123L complex contained a total of 312,693 particles in the closed state. These particles were sorted into five distinct classes of 3.2 Å resolution. The CCT-PhLP1-R269E/R311E complex was processed as described for the other mutants except the unbinned box size was 512 pixels. The final dataset contained a total of 232,800 particles in the closed state. After classification the particles were sorted into two classes of 3.0 Å resolution and one class of 3.4 Å resolution.

For comparison, the same image processing was done for the WT CCT-PhLP1-Gβ_5_ complex using a similar number of particles as the mutants. This dataset followed a similar workflow as previously described for the WT complex (37). From the dataset previously published (EMPIAR-12086) 25,455 micrographs were motion corrected, CTF estimated, and curated with a 5 Å cutoff. After particle selection with a blob picker, extraction with a box size of 300 pixels, and iterative 2D classification, 2,393,120 particles were used for Ab initio reconstruction and heterogenous refinement. 1,868,630 closed particles were randomly divided into subsets containing 200,000 or 300,000 particles.

When the dataset was divided into subsets of 300,000 particles, six subsets contained 300,000 particles and one subsect contained 68,630 particles. One of the 300,000 particle subsets was selected for focused 3DVA, with a focus mask over the substrate folding chamber, and non-uniform refinement. The component showing the greatest degree of continuous motion was selected for further focused 3D classification using a mask over half of the chamber containing Gβ_5._ After non-uniform refinement, this workflow produced six reconstructions, with one reconstruction containing 54,905 particles and a fully folded Gβ_5._

When the dataset was divided into subsets of 200,000 particles, 9 subsets contained 200,000 particles and one class contained 68,630 particles. One of the 200,000 subsets was selected for focused 3DVA, non-uniform refinement, focused classification, and a final round of non-uniform refinement. This workflow resulted in four reconstructions. One of these reconstructions contained 52,731 particles and a fully folded Gβ_5._

### Model refinement

UCSF ChimeraX (version 1.8) (62) was used to rigid body fit previous models of CCT-PhLP1-Gβ_5_ into the density for all mutant classes. PDB: 8SFE (37) was used as the starting model for CCT and PhLP1 while PDB: 8SGL (37) was used as the starting model for Gβ_5_. Models were iteratively real-space refined using Phenix v1.20.1 (63) with ADP, AlF_3_, H_2_O and Mg^2+^ restraint files and manually adjusted in Coot v0.9.8.5 (64). In parallel, all classes were also refined with the 8SGL model rotated into the alternate orientation of Gβ_5._

The orientation of blades in each reconstruction was determined by rigid body fitting a model of fully folded Gβ_5_ (PDB: 8SGL) into the classes of each mutant. For R269E, the density between CCT5 and Gβ_5_ at the mutation site indicated a potential alternate orientation of Gβ_5_ within CCT. All blades of the 8SGL model were rigid body fit into the Gβ_5_ R269E density. Only one other blade contained a positively charged residue on the outside of the blade that was in register with the density. Models were refined for both orientations, and the fit of residues in both the WT and alternate orientations into the reconstruction density was visually assessed.

### Sequence alignment

The sequence of Gβ_5_ was truncated to correspond with each of the seven blades and were aligned using Jalview (65) (version 2.11.4.1). ConSurf was used to analyze the Gβ_5_ sequence conservation, and blade sequences were colored according to their level of conservation as calculated with ConSurf.

### Clash analysis

Prediction of steric clashes for the G257E and S123L mutants with neighboring blades were performed by opening a model of fully folded Gβ_5_ (PDB:8SGL) with UCSF ChimeraX (version 1.8). The mutations were made using the command line based swapaa command utilizing the Dunbrack rotamer library (66). Rotamers were then chosen based on the lowest clash score. Clashes were predicted based on a van der Waals overlap cutoff of ≥ 0.6 Å with an hbond allowance of 0.4Å. Prediction of steric clashes between R269E blade 3 with CCT4 and blade 6 with CCT5 was performed by opening a model of fully folded Gβ_5_ (PDB:8SGL) with UCSF ChimeraX. The model for Gβ_5_ was rotated into the alternate conformation, and clashes were predicted as described above.

## Data availability

Cryo-EM reconstructions and atomic models have been deposited in the Electron Microscopy Data Bank (EMDB) and Protein Data Bank (PDB), respectively. Accession numbers for deposited structures are listed in *SI Appendix* Tables 2-4. Any additional information required to reanalyze the data reported in this paper is available from the corresponding authors upon reasonable request.

## Acknowledgments

This work was supported by NIH grant R01 EY036925 to P.S.S. and B.M.W, NIH grant R35 GM133772 to P.S.S., NIH grant R01 EY012287 to B.M.W., NIH training grant T32 EY02434 to M.I.S, NIH Predoctoral Fellowship F31 GM150221 to D.C.M, and the BYU Simmons Center for Cancer Research (M.I.S). We thank David Belnap and the University of

Utah Arnold and Mabel Beckman Center for Cryo-EM as well as the BYU Electron Microscopy Facility for cryo-EM support. We also thank the University of Utah Center for High Performance Computing and the BYU Office of Research Computing for computational support. Finally, we acknowledge support from the BYU Department of Chemistry and Biochemistry Confocal Microscopy facility.

## Author Contributions

Conceptualization, B.M.W. and P.S.S.; Methodology, B.M.W., P.S.S., M.I.S., D.C.M., and S.M.; Investigation, M.I.S., D.C.M., R.A.N., C.A.J., J.N.R.-T., A.M.W.,

R.L.B., C.S.S., S.L.C., L.J.M., and P.S.S.; Data Curation, D.C.M., R.A.N., and P.S.S.; Writing,

M.I.S., D.C.M., B.M.W., and P.S.S.; Visualization, D.C.M., R.A.N., M.I.S., B.M.W., and P.S.S.;

Funding Acquisition, B.M.W. and P.S.S.; Supervision, B.M.W. and P.S.S.

