## Supplemental Material for "Disease-causing mutations in the G protein β5 β-propeller disrupt its chaperonin-mediated folding trajectory"

### SI Appendix Figures and Tables

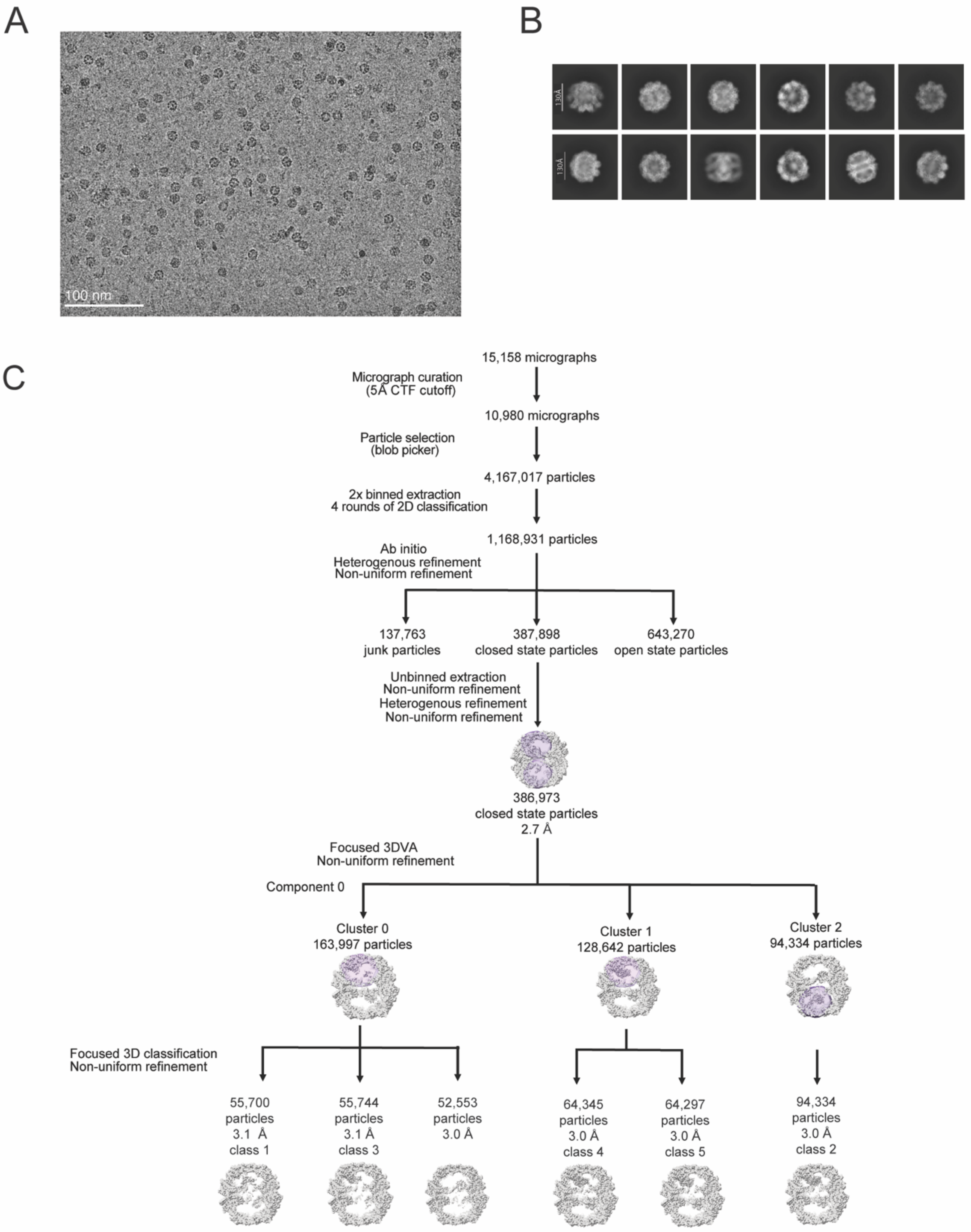


**Figure S1. Image processing workflow of CCT-PhLP1-G$\beta_5$ R269E.** (A) A representative cryo-EM micrograph. (B) Representative 2D class averages. Scale bar represents 130 Å. (C) Workflow of data processing for CCT-PhLP1-G$\beta_5$ R269E. Masks used for focused 3DVA and focused 3D classification are shown in purple.

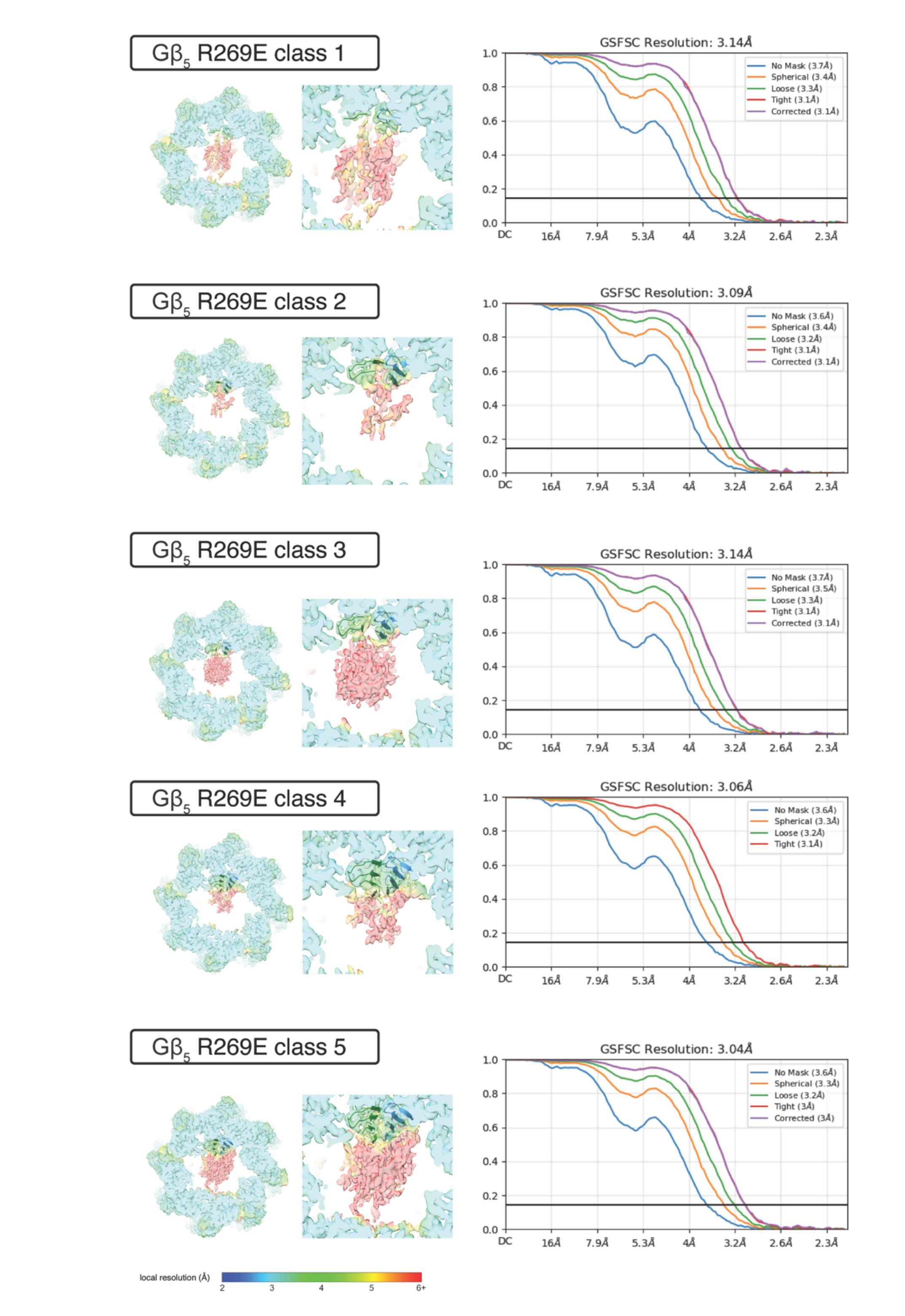


**Figure S2. Reconstructions of CCT-PhLP1-Gβ$_5$ R269E.** Sections show top views of reconstructions for each folding intermediate with Gβ$_5$-focused insets colored by local resolution (left) as well as Fourier shell correlation (FSC) curves (right).

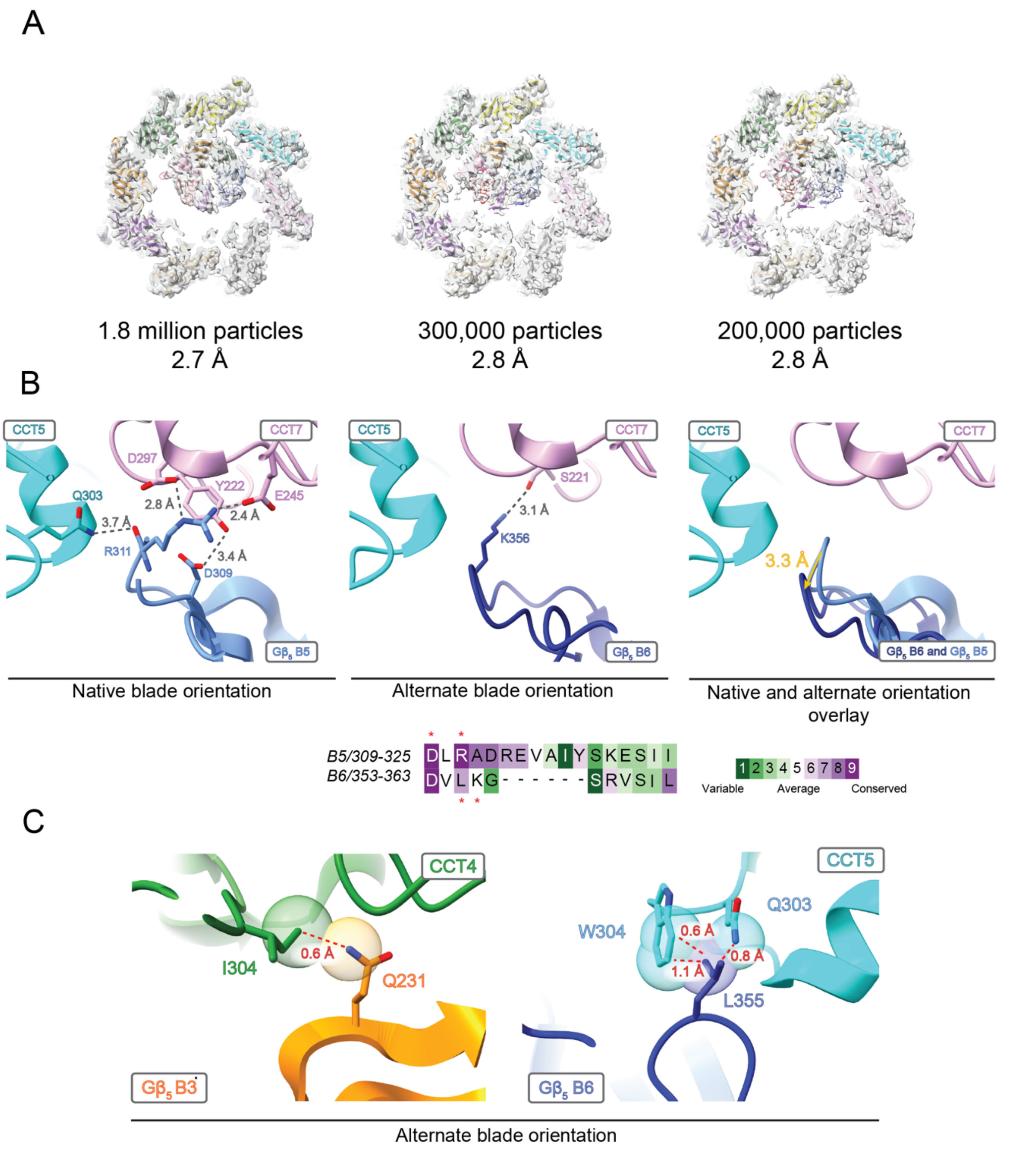


**Figure S3. Structures of CCT-PhLP1-Gβ$_5$ WT and predicted clashes of Gβ$_5$ R269E.** (A) (Left) WT CCT-PhLP1-Gβ$_5$ processed with a full set of 1.8 million particles, (Middle) a random subset of 300,000 particles, and (Right) a random subset of 200,000 particles, all showing a closed β-propeller, for comparison with the Gβ$_5$ mutant structures. (B) (Left) Interactions between Gβ$_5$ blade 5 (B5) and CCT subunits in the native orientation and (Middle) between blade 6 (B6) and CCT subunits in the alternate blade orientation. (Right) Displacement of Gβ$_5$ blade 6 away from CCT7 in the alternate orientation (dark blue) compared to blade 5 in the native orientation (light blue). (Below) Alignment and conservation analysis of the outer strands of blade 5 and blade 6 (B6). (C) (Left) Predicted clashes between Gβ$_5$ R269E blade 3 (B3) and CCT4 and (Right) predicted clashes between CCT5 and Gβ$_5$ R269E blade 6 in the alternate orientation. Red dashes and labels indicate van der Waals overlap distances.

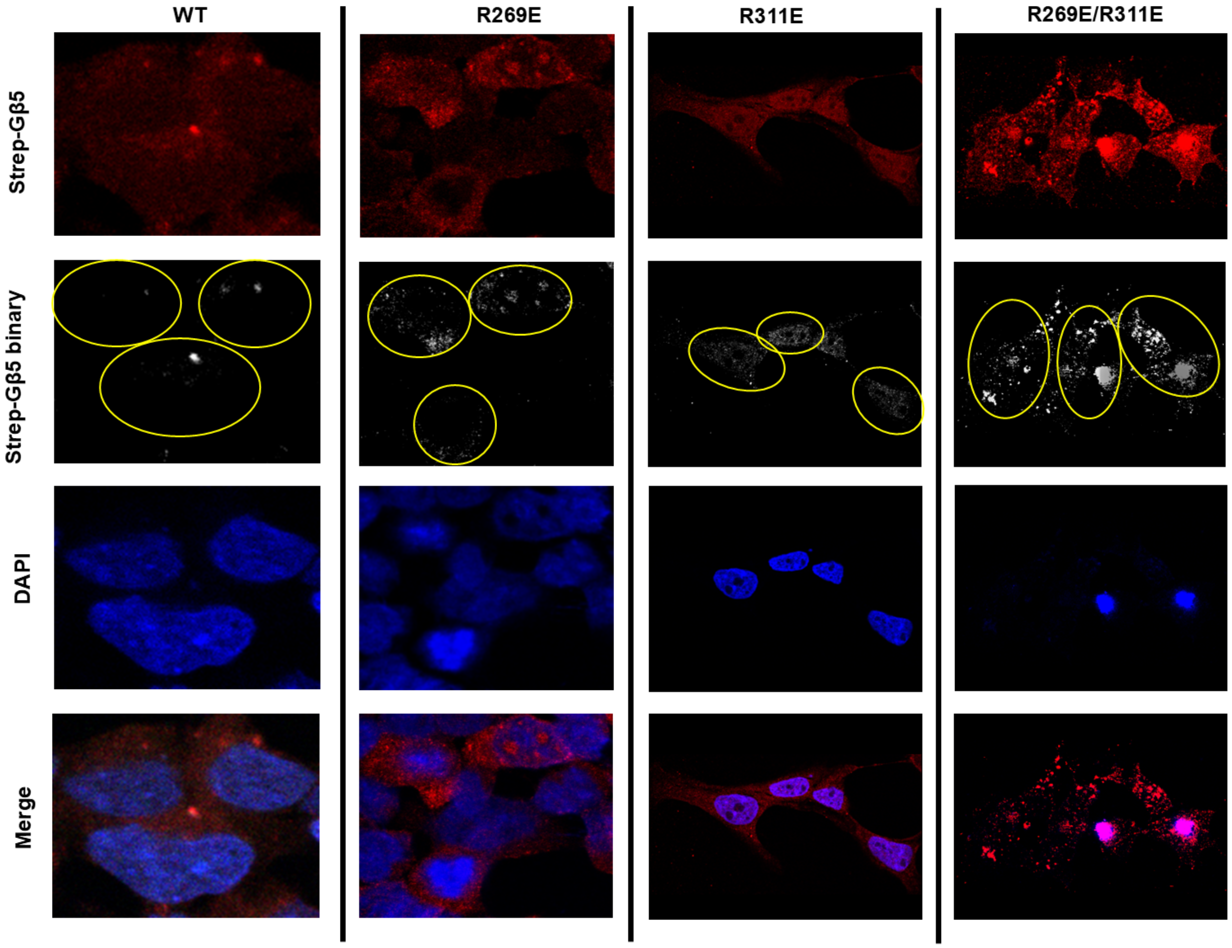


**Figure S4. G$\beta_5$ R269E, R311E, and R269E/R311E immunolocalization.** HEK-293T cells were transfected with the indicated Strep-G$\beta_5$ mutants and immunolocalized with an anti-Strep antibody (red). The images were transformed into binary format, and the puncta were quantified using the Fiji software as described in Methods. Cells used in the quantification are outlined in yellow. Nuclei were stained with DAPI (blue). Representative images are shown.

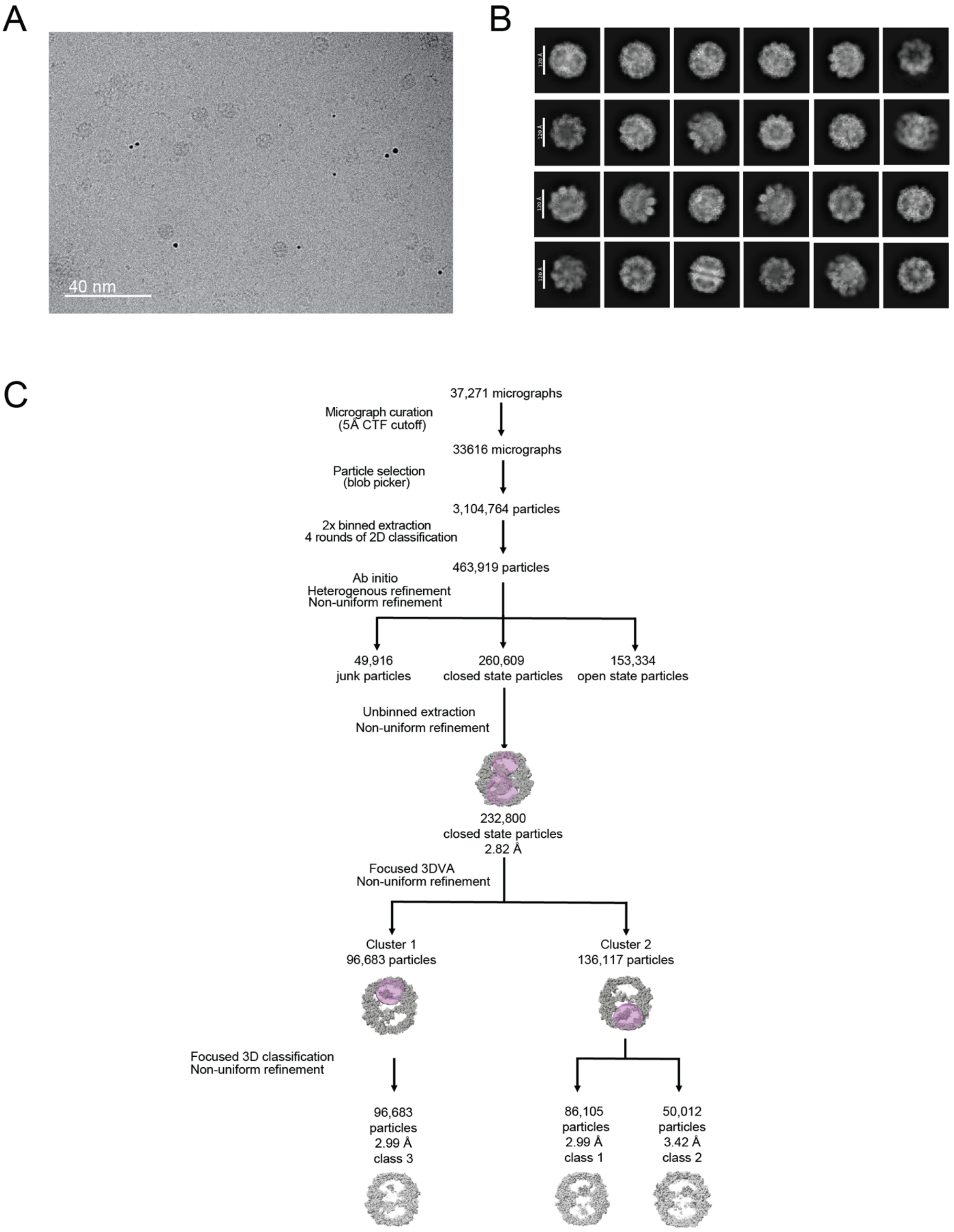


**Figure S5. Image processing workflow of CCT-PhLP1-G$\beta_5$ R269E/R311E.** (A) A representative cryo-EM micrograph. (B) Representative 2D class averages. Scale bar represents 120 Å. (C) Workflow of data processing for CCT-PhLP1-G$\beta_5$ R269E/R311E. Masks used for focused 3DVA and focused 3D classification are shown in purple.

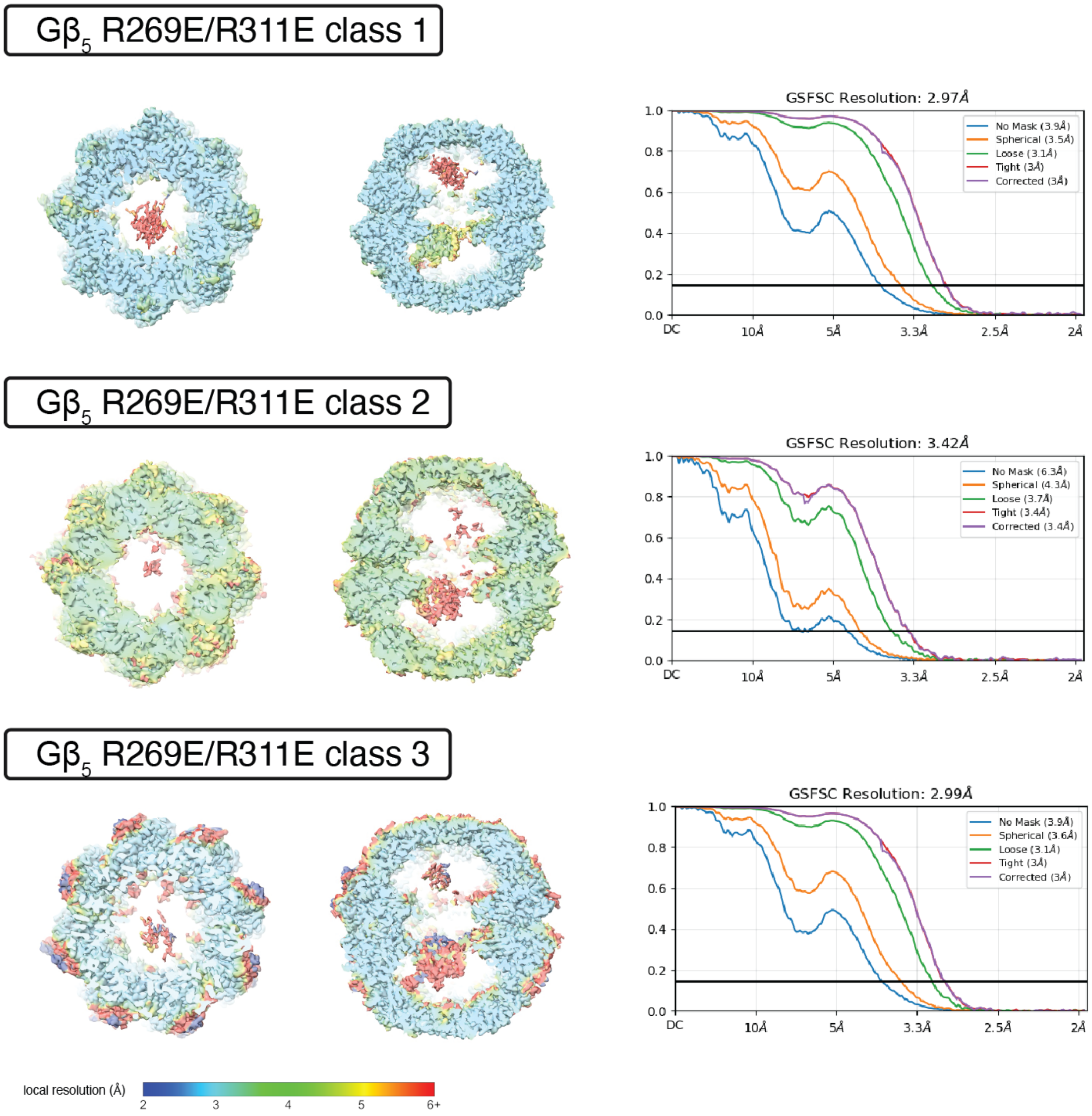


**Figure S6. Reconstructions of CCT-PhLP1-Gβ5 R269E/R311E.** Sections show top and side views of reconstructions for each folding intermediate colored by local resolution (left) as well as Fourier shell correlation (FSC) curves (right).

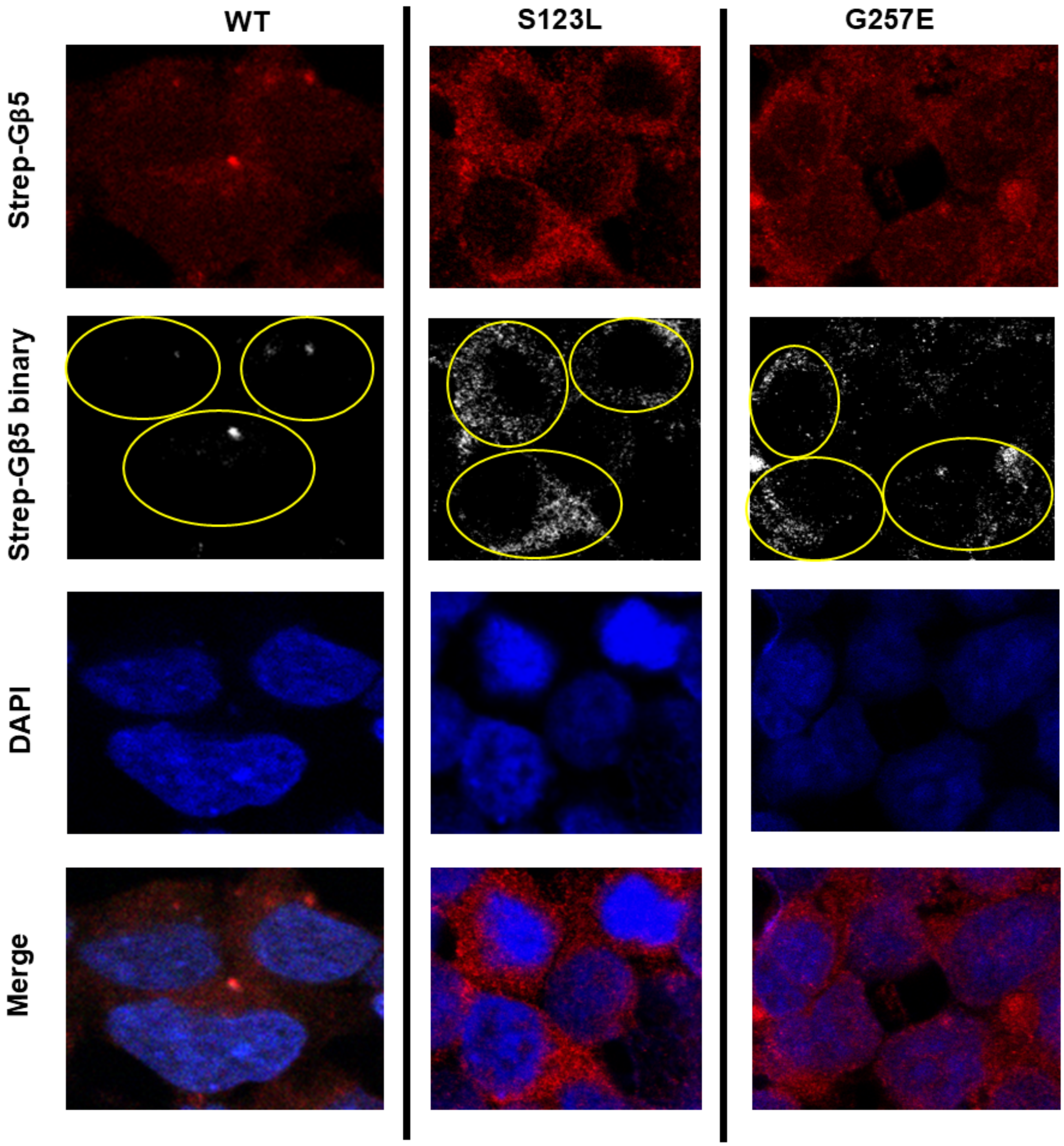


**Figure S7. G$\beta_5$ S123L and G257E immunolocalization.** HEK-293T cells were transfected with the indicated Strep-G$\beta_5$ mutants and immunolocalized with an anti-Strep antibody (red). The images were transformed into binary format, and the puncta were quantified using the Fiji software as described in Methods. Cells used in the quantification are outlined in yellow. Nuclei were stained with DAPI (blue).

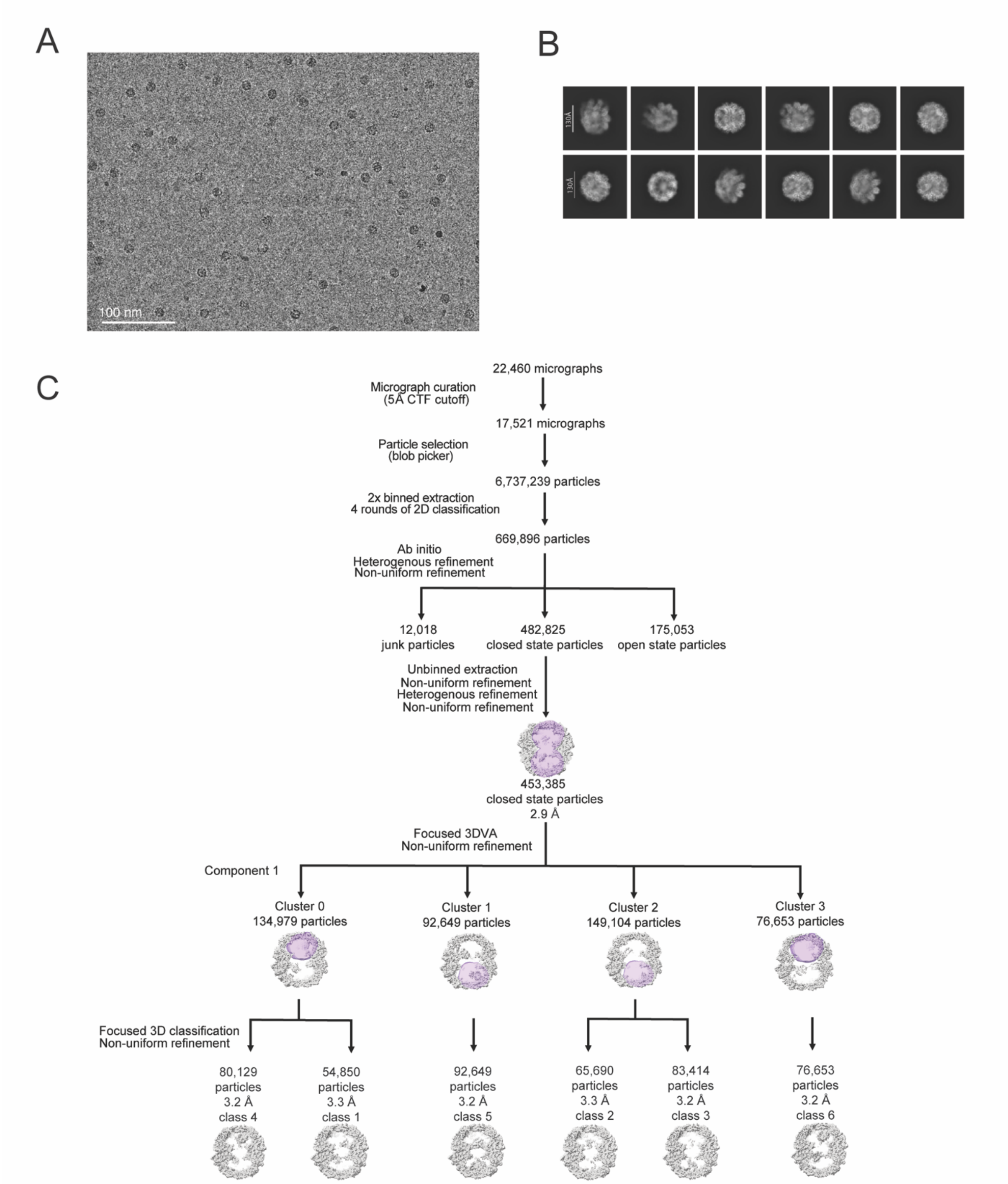


**Figure S8. Image processing workflow of CCT-PhLP1-G$\beta_5$ G257E.** (A) A representative cryo-EM micrograph. (B) Representative 2D class averages. Scale bar represents 130 Å. (C) Workflow of data processing for CCT-PhLP1-G$\beta_5$ G257E. Masks used for focused 3DVA and focused 3D classification are shown in purple.

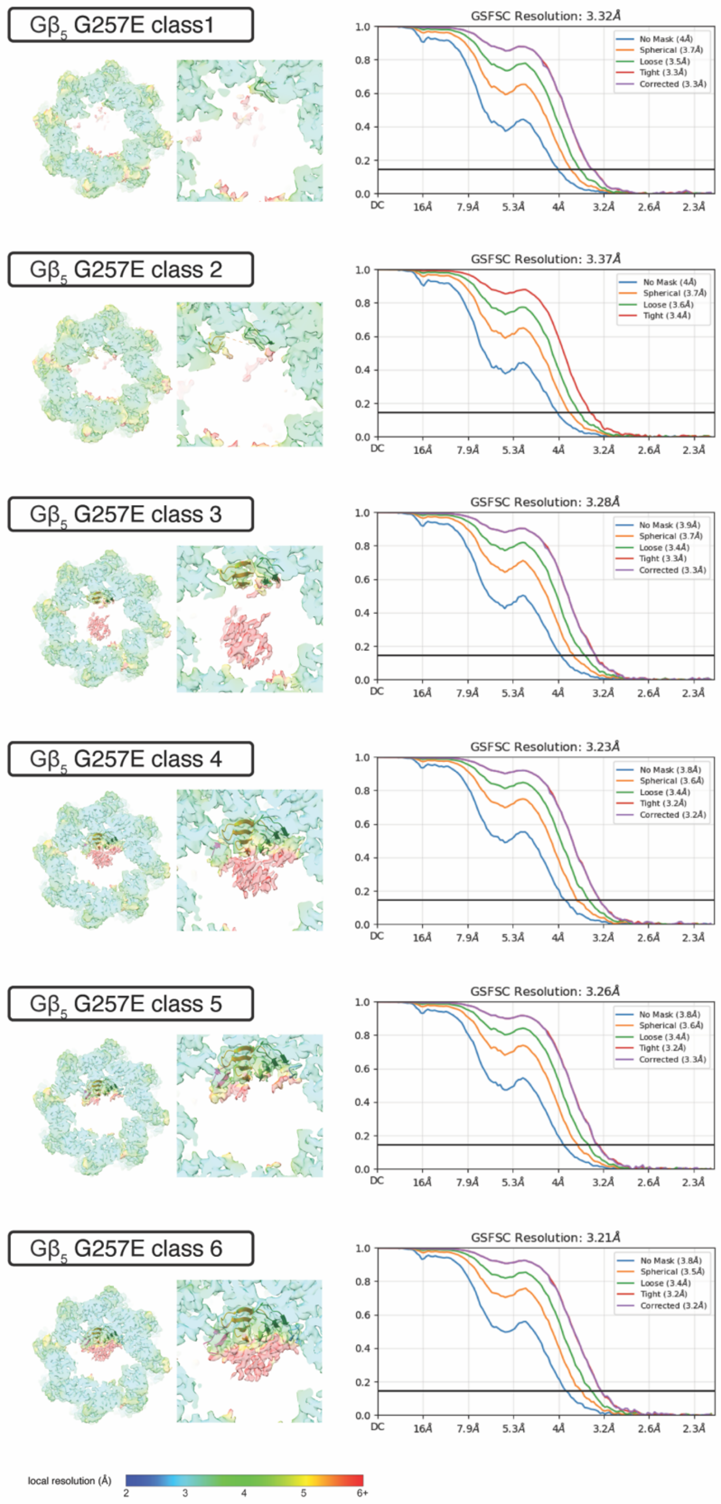


**Figure S9. Reconstructions of CCT-PhLP1-Gβ5 G257E.** Sections show top views of reconstructions for each folding intermediate with Gβ5-focused insets colored by local resolution (left) as well as Fourier shell correlation (FSC) curves (right).

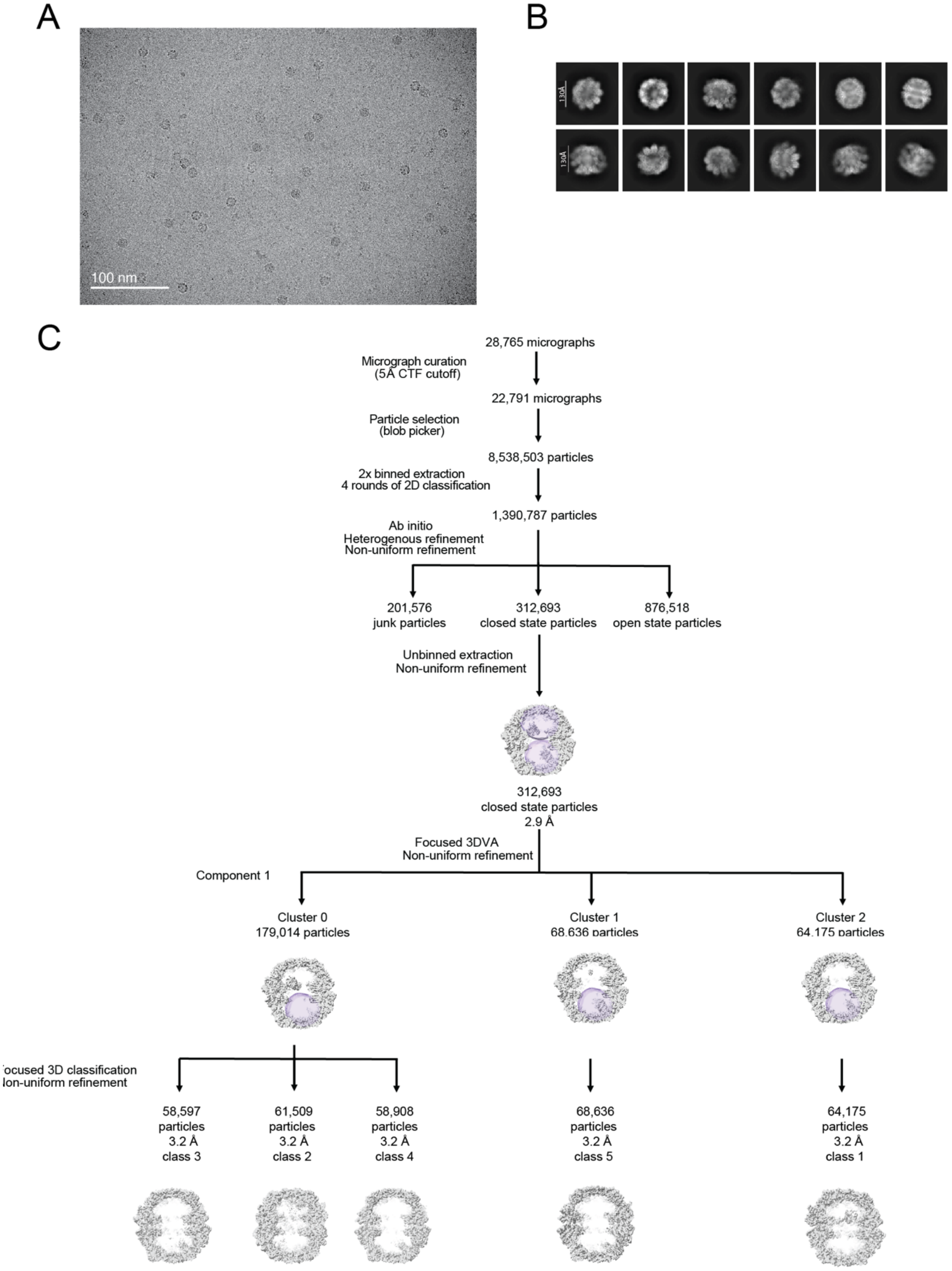


**Figure S10. Image processing workflow of CCT-PhLP1-G$\beta_5$ S123L.** (A) A representative cryo-EM micrograph. (B) Representative 2D class averages. Scale bar represents 130 Å. (C) Workflow of data processing for CCT-PhLP1-G$\beta_5$ S123L. Masks used for focused 3DVA and focused 3D classification are shown in transparent purple.

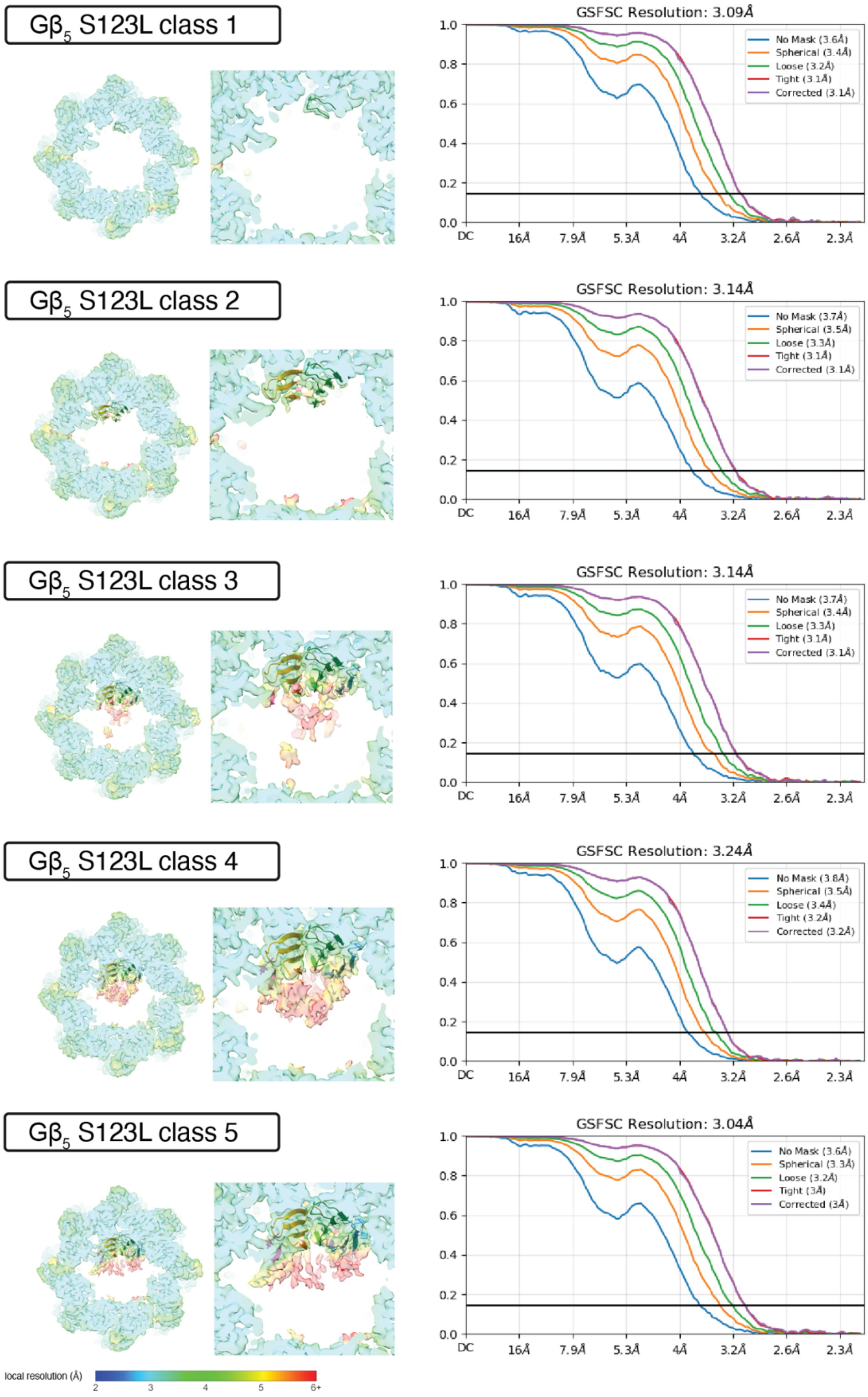


**Figure S11. Reconstructions of CCT-PhLP1-Gβ5 S123L.** Sections show top views of reconstructions for each folding intermediate with Gβ5-focused insets colored by local resolution (left) as well as Fourier shell correlation (FSC) curves (right).

**Supplemental Table S1. Cryo-EM Data Collection Parameters (1/2)**

| **Data collection** | | | | | | | | | |
|---|---|---|---|---|---|---|---|---|---|
| Microscope | Titan Krios G3 | | | | | | | | |
| Voltage (kV) | 300 | | | | | | | | |
| Detector | Gatan K3 | | | | | | | | |
| Energy Filter | BioQuantum | | | | | | | | |
| Energy Slit (eV) | 10 | | | | | | | | |
| Data collection software | SerialEM | | | | | | | | |
| Nominal magnification | 81,000x | | | | | | | | |
| Dataset | G$\beta_5$ R269E | | | G$\beta_5$ G257E | | G$\beta_5$ S123L | | | |
| Dose rate ($e^-/Å^2$/second) | 16.8 | 13.5 | 13.2 | 16.8 | 17.0 | 18.2 | 17.5 | 17.2 | 16.9 |
| Total number of frames | 40 | | | | | | | | |
| Total electron exposure ($e^-/Å^2$) | 42.1 | 33.7 | 33.1 | 42.0 | 42.6 | 45.4 | 43.7 | 43.0 | 42.2 |
| Defocus range (µm) | -0.8 to -1.2 | | | | | | | | |
| Pixel size (Å) | 1.058 | | | | | | | | |
| **Data processing** | | | | | | | | | |
| Number of micrographs | 15,158 | | | 22,460 | | 28,765 | | | |
| Final number of closed CCT particles | 386,973 | | | 453,385 | | 312,693 | | | |
| Symmetry imposed | C1 | | | | | | | | |

**Supplemental Table S1. Cryo-EM Data Collection Parameters (2/2)**

| **Data collection** | | | |
|---|---|---|---|
| Microscope | Titan Krios G4 | | |
| Voltage (kV) | 300 | | |
| Detector | Falcon 4 | | |
| Energy Filter | Selectris X | | |
| Energy Slit (eV) | 10 | | |
| Data collection software | EPU | | |
| Nominal magnification | 130,000x | | |
| Dataset | G$\beta_5$ R269E/R311E | | |
| Dose rate ($e^-/Å^2$/second) | 9.34 | 7.68 | 9.52 |
| Total number of frames | Not Relevant (Collected .eer) | | |
| Total electron exposure ($e^-/Å^2$) | 25.95 | 33.27 | 26.46 |
| Defocus range (µm) | -0.6 to -1.6 | | |
| Pixel size (Å) | 0.95 | | |
| **Data processing** | | | |
| Number of micrographs | 37,271 | | |
| Final number of closed CCT particles | 232,800 | | |
| Symmetry imposed | C1 | | |

**Supplemental Table S2. 3D reconstruction and model refinement statistics (1/3)**

| **Structure** | R269E class 1 | R269E class 2 | R269E class 3 | R269E class 4 | R269E class 5 |
|---|---|---|---|---|---|
| EM Databank Accession ID | EMD-49639 | EMD-49634 | EMD-49732 | EMD-49733 | EMD-49604 |
| Protein Data Bank Accession ID | 9NQ1 | 9NPW | 9NRG | 9NRH | 9NOQ |
| **Symmetry imposed** | | | | | |
| 3D classification | C1 | | | | |
| 3D refinement | C1 | | | | |
| Final particles | 55,700 | 94,334 | 55,744 | 64,345 | 64,297 |
| **Map resolution (Å)** | | | | | |
| FSC 0.143 (unmasked) | 3.7 | 3.6 | 3.7 | 3.6 | 3.6 |
| FSC 0.143 (masked, corrected) | 3.1 | 3.1 | 3.1 | 3.1 | 3.0 |
| **Model Refinement** | | | | | |
| Initial model used for CCT (PDB code) | 8SH9 | | | | |
| Initial model used for G$\beta_5$ (PDB code) | 8SGL | | | | |
| Initial model used for PhLP1 (PDB code) | 8SH9 | | | | |
| Map correlation coefficient | 0.88 | 0.88 | 0.88 | 0.87 | 0.89 |
| **Model composition** | | | | | |
| Non-hydrogen atoms | 65,252 | 66,098 | 65,402 | 67,257 | 67,278 |
| Protein residues | 8,476 | 8,588 | 8,496 | 8,730 | 8,733 |
| Ligands (ADP, $AlF_x$, $Mg^{2+}$) | 16 | | | | |
| **R.m.s. deviations** | | | | | |
| Bond lengths (Å) | 0.005 | 0.004 | 0.005 | 0.006 | 0.004 |
| Bond angles (˚) | 1.026 | 0.597 | 0.680 | 0.705 | 0.625 |
| **Validation** | | | | | |
| MolProbity score | 1.69 | 1.60 | 1.69 | 1.69 | 1.67 |
| Clashscore | 9.18 | 7.78 | 9.04 | 8.86 | 9.20 |
| **Ramachandran plot** | | | | | |
| Favored (%) | 96.76 | 97.01 | 96.75 | 96.62 | 96.91 |
| Allowed (%) | 3.24 | 2.96 | 3.24 | 3.36 | 3.08 |
| Outliers (%) | 0 | 0.02 | 0.01 | 0.01 | 0.01 |
| C-beta deviations (0.25 Å) | 0 | 0 | 0 | 0 | 0 |
| CaBLAM outliers (%) | 1.70 | 1.77 | 1.75 | 1.80 | 1.91 |
| EMRinger Score | 2.17 | 2.33 | 2.26 | 2.36 | 2.34 |

**Supplemental Table S2, continued. 3D reconstruction and model refinement statistics (2/3)**

| **Structure** | G257E class 1 | G257E class 2 | G257E class 3 | G257E class 4 | G257E class 5 | G257E class 6 |
|---|---|---|---|---|---|---|
| EM Databank Accession ID | EMD-49709 | EMD-49729 | EMD-49710 | EMD-49642 | EMD-49730 | EMD-49731 |
| Protein Data Bank Accession ID | 9NR1 | 9NRD | 9NR4 | 9NQ6 | 9NRE | 9NRF |
| **Symmetry imposed** | | | | | | |
| 3D classification | C1 | | | | | |
| 3D refinement | C1 | | | | | |
| Final particles | 54,850 | 65,690 | 83,414 | 80,129 | 92,649 | 76,653 |
| **Map resolution (Å)** | | | | | | |
| FSC 0.143 (unmasked) | 4.0 | 4.0 | 3.9 | 3.8 | 3.8 | 3.8 |
| FSC 0.143 (masked, corrected) | 3.3 | 3.4 | 3.3 | 3.3 | 3.2 | 3.2 |
| **Model Refinement** | | | | | | |
| Initial model used for CCT (PDB code) | 8SH9 | | | | | |
| Initial model used for G$\beta_5$ (PDB code) | 8SGL | | | | | |
| Initial model used for PhLP1 (PDB code) | 8SH9 | | | | | |
| Map correlation coefficient | 0.87 | 0.88 | 0.86 | 0.87 | 0.87 | 0.88 |
| **Model composition** | | | | | | |
| Non-hydrogen atoms | 65,214 | 65,279 | 65,421 | 67,378 | 65,870 | 67,578 |
| Protein residues | 8,472 | 8,481 | 8,499 | 8,753 | 8,560 | 8,776 |
| Ligands (ADP, $AlF_x$, $Mg^{2+}$) | 16 | | | | | |
| **R.m.s. deviations** | | | | | | |
| Bond lengths (Å) | 0.004 | 0.005 | 0.005 | 0.006 | 0.005 | 0.005 |
| Bond angles (˚) | 0.568 | 0.606 | 0.615 | 0.731 | 0.647 | 0.605 |
| **Validation** | | | | | | |
| MolProbity score | 1.64 | 1.74 | 1.70 | 1.75 | 1.69 | 1.66 |
| Clashscore | 9.10 | 10.18 | 9.33 | 10.13 | 8.97 | 8.88 |
| **Ramachandran plot** | | | | | | |
| Favored (%) | 97.14 | 96.71 | 96.71 | 96.51 | 96.64 | 96.87 |
| Allowed (%) | 2.85 | 3.29 | 3.29 | 3.48 | 3.35 | 3.13 |
| Outliers (%) | 0.01 | 0 | 0 | 0.01 | 0.01 | 0 |
| C-beta deviations (0.25 Å) | 0 | 0 | 0 | 0 | 0 | 0 |
| CaBLAM outliers (%) | 1.73 | 1.70 | 1.73 | 1.81 | 1.78 | 1.90 |
| EMRinger Score | 1.71 | 1.59 | 1.90 | 2.05 | 2.17 | 2.03 |

**Supplemental Table S2, continued. 3D reconstruction and model refinement statistics (3/3)**

| **Structure** | S123L class 1 | S123L class 2 | S123L class 3 | S123L class 4 | S123L class 5 |
|---|---|---|---|---|---|
| EM Databank Accession ID | EMD-72144 | EMD-72107 | EMD-72098 | EMD-72106 | EMD-72096 |
| Protein Data Bank Accession ID | 9Q1X | 9Q0V | 9Q0G | 9Q0U | 9Q0E |
| **Symmetry imposed** | | | | | |
| 3D classification | C1 | | | | |
| 3D refinement | C1 | | | | |
| Final particles | 64,175 | 61,509 | 58,597 | 58,908 | 68,636 |
| **Map resolution (Å)** | | | | | |
| FSC 0.143 (unmasked) | 3.7 | 3.8 | 3.8 | 3.8 | 3.7 |
| FSC 0.143 (masked, corrected) | 3.2 | 3.2 | 3.2 | 3.2 | 3.2 |
| **Model Refinement** | | | | | |
| Initial model used for CCT (PDB code) | 8SH9 | | | | |
| Initial model used for $G\beta_5$ (PDB code) | 8SGL | | | | |
| Initial model used for PhLP1 (PDB code) | 8SH9 | | | | |
| Map correlation coefficient | 0.88 | 0.88 | 0.89 | 0.88 | 0.88 |
| **Model composition** | | | | | |
| Non-hydrogen atoms | 66,320 | 65,456 | 65,802 | 65,847 | 66,320 |
| Protein residues | 8,611 | 8,503 | 8,551 | 8,556 | 8,611 |
| Ligands (ADP, $AlF_x$, $Mg^{2+}$) | 16 | | | | |
| **R.m.s. deviations** | | | | | |
| Bond lengths (Å) | 0.005 | 0.005 | 0.005 | 0.005 | 0.005 |
| Bond angles (˚) | 0.954 | 0.960 | 0.977 | 0.961 | 0.954 |
| **Validation** | | | | | |
| MolProbity score | 1.57 | 1.60 | 1.66 | 1.61 | 1.57 |
| Clashscore | 7.91 | 8.07 | 8.60 | 8.24 | 7.91 |
| **Ramachandran plot** | | | | | |
| Favored (%) | 97.23 | 97.09 | 96.83 | 97.09 | 97.23 |
| Allowed (%) | 2.77 | 2.91 | 3.17 | 2.91 | 2.77 |
| Outliers (%) | 0 | 0 | 0 | 0 | 0 |
| C-beta deviations (0.25 Å) | 0 | 0 | 0 | 0 | 0 |
| CaBLAM outliers (%) | 1.73 | 1.60 | 1.87 | 1.82 | 1.73 |
| EMRinger Score | 1.91 | 2.06 | 2.07 | 2.15 | 1.91 |

**Supplemental Table S3. R269E/R311E 3D reconstruction statistics**

| **Structure** | R269E/R311E class 1 | R269E/R311E class 2 | R269E/R311E class 3 |
|---|---|---|---|
| EM Databank Accession ID | Pending | Pending | Pending |
| **Symmetry imposed** | | | |
| 3D classification | | C1 | |
| 3D refinement | | C1 | |
| Final particles | 86,105 | 50,012 | 96.683 |
| **Map resolution (Å)** | | | |
| FSC 0.143 (unmasked) | 3.9 | 6.3 | 3.9 |
| FSC 0.143 (masked, corrected) | 3.0 | 3.4 | 3.0 |
